# Structural basis of Rgg protein binding to their regulatory pheromones and target DNA promoters

**DOI:** 10.1101/2020.04.30.069369

**Authors:** Glenn C. Capodagli, Kaitlyn M. Tylor, Jason T. Kaelber, Vasileios I. Petrou, Michael J. Federle, Matthew B. Neiditch

**Author notes:** Corresponding Authors: Matthew B. Neiditch, Michael J. Federle, Vasileios I. Petrou. These authors contributed equally to this work (Glenn C. Capodagli, Kaitlyn M. Tylor). Author Contributions G.C.C., K.M.T., J.T.K., V.I.P, M.J.F., and M.B.N. designed research, performed research, contributed new reagents/analytic tools, analyzed data, and wrote the paper.

## Abstract

Rgg family proteins, such as Rgg2 and Rgg3, have emerged as primary quorum-sensing regulated transcription factors in *Streptococcus* species, controlling virulence, antimicrobial resistance, and biofilm formation. Rgg2 and Rgg3 function is regulated by their interaction with oligopeptide quorum-sensing signals called short hydrophobic peptides (SHPs). The molecular basis of Rgg-SHP and Rgg-target DNA promoter specificity was unknown. To close this gap, we determined the cryo-EM structure of *Streptococcus thermophilus* Rgg3 bound to its quorum-sensing signal, SHP3, and the X-ray crystal structure of Rgg3 alone. Comparison of these structures to that of an Rgg in complex with cyclosporin A (CsA), an inhibitor of SHP-induced Rgg activity, reveals the molecular basis of CsA function. Furthermore, to determine how Rgg proteins recognize DNA promoters, we determined X-ray crystal structures of both *S. dysgalactiae* Rgg2 and *S. thermophilus* Rgg3 in complex with their target DNA promoters. The physiological importance of the observed Rgg-DNA interactions was dissected using *in vivo* genetic experiments and *in vitro* biochemical assays. Based on these structure-function studies, we present a revised unifying model of Rgg regulatory interplay. In contrast to existing models, where Rgg2 proteins are transcriptional activators and Rgg3 proteins are transcriptional repressors, we propose that both are capable of transcriptional activation. However, when Rgg proteins with different activation requirements compete for the same DNA promoters, those with more stringent activation requirements function as repressors by blocking promoter access of the SHP-bound conformationally active Rgg proteins. While a similar gene expression regulatory scenario has not been previously described, in all likelihood it is not unique to streptococci.

**Significance Statement:** Secreted peptide pheromones regulate critical biological processes in Gram-positive bacteria. In streptococci such as the human pathogen *S. pyogenes*, oligopeptide pheromones, like the short hydrophobic peptides (SHPs), regulate virulence, antimicrobial resistance, and biofilm formation. SHPs directly regulate the activity of transcription factors called Rgg2 and Rgg3. We present the cryo-EM structure of Rgg3 in complex with SHP3, as well as X-ray crystal structures of Rgg2 bound to target promoter DNA, Rgg3 bound to target promoter DNA, and Rgg3 alone. Based on the cryo-EM, X-ray crystallographic, biochemical, and genetic studies presented here, we provide not only detailed mechanistic insight into the molecular basis of Rgg3-SHP3, Rgg2-DNA, and Rgg3-DNA binding specificity, but also a new model of transcription factor regulatory interplay.

## Introduction

Gram-positive bacteria use oligopeptides as signals for cell-cell communication, a process known as quorum sensing (1). The oligopeptide signals (also known as pheromones or autoinducers) are synthesized as pro-peptides, usually containing an N-terminal secretion signal. The pro-peptides are proteolytically processed to their mature form and secreted. The pheromones then bind to and regulate the activity of a membrane-bound receptor, or otherwise bind to an oligopeptide permease that transports the pheromone into the cytoplasm. Here, the pheromones bind receptors called RRNPP proteins, named after the family member archetypes <u>R</u>ap, <u>R</u>gg, <u>N</u>prR, <u>P</u>lcR, and <u>P</u>rgX.

The primary family of quorum-sensing receptors in *Streptococcus* species are the Rgg proteins, which, for example, regulate virulence, antimicrobial resistance, and competence-related genes in the human pathogen *Streptococcus pyogenes* (Group A *Streptococcus*)(2-7). Rgg proteins are DNA-binding transcription factors, whose activity is regulated by oligopeptide pheromones such as the short hydrophobic peptides (SHPs) (1, 8-10).

It was previously reported that nearly all streptococci encode an *rgg* gene, with some species containing multiple paralogs (11). Two families of Rgg proteins, Rgg2 and Rgg3, have been described based on their sequence similarity (9). Following genetic and biochemical studies in *S. pyogenes*, it was proposed that Rgg2 proteins are transcriptional activators while Rgg3 proteins are transcriptional repressors (9). The SHPs that target Rgg2 and Rgg3 proteins are called SHP2 and SHP3, respectively, although there can be crosstalk, e.g., *S. pyogenes* SHP2 and SHP3 have similar amino acid sequences and bind to both Rgg2_sp_ and Rgg3_Sp_ (5). Furthermore, the mature SHP2 and SHP3 peptides have been categorized as belonging to one of three groups (10). Group I and II SHPs contain an N-terminal aspartate or glutamate, respectively, and they are typically eight or nine amino acids long. Group I and II *shp* genes are located in close proximity to the genes encoding their cognate Rgg target proteins from which they are divergently transcribed. Group III SHPs contain an N-terminal aspartate or glutamate, but they are encoded by genes overlapping the ends of the *rgg* genes from which they are convergently transcribed. The acidic residues at the N-terminus of the SHPs are required for their activity, but the reason for this requirement was unknown (5, 8, 12). The structural basis of SHP binding to Rgg2 and Rgg3 proteins, as well as the structural basis of Rgg binding to DNA, were also unknown.

In previous studies we determined the X-ray crystal structures of *Streptococcus dysgalactiae* Rgg2 (Rgg2_Sd_) alone and in complex with cyclosporin A (CsA), which we proposed is a competitive inhibitor of SHP2 and SHP3 binding (13). These structural studies along with complementary *in vitro* biochemical and *in vivo* genetic analyses showed that Rgg2 and Rgg3 proteins contain an N-terminal XRE-family DNA-binding domain and a C-terminal tetratricopeptide-like SHP2 and SHP3-binding domain (called the repeat domain). These studies provided insight into the Rgg2 structure only, and not Rgg3. Most importantly, they did not reveal how Rgg2 or Rgg3 proteins function to specifically recognize their cognate SHPs or target DNA promoters. In addition, because we did not understand how SHPs bind to Rgg proteins, it was not clear how CsA functions to inhibit their activity.

To address these gaps we determined the cryo-EM structure of *Streptococcus thermophilus* Rgg3 (Rgg3_St_) in complex with SHP3, the X-ray crystal structures of Rgg3_St_ alone, the X-ray crystal structure of Rgg2_Sd_ in complex with its target Rgg-box DNA, and the X-ray crystal structure of Rgg3_St_ in complex with its target Rgg-box. Based on the cryo-EM, X-ray crystallographic, biochemical, and genetic studies presented here, we provide not only detailed mechanistic insight into Rgg-SHP, Rgg2-DNA, and Rgg3-DNA interactions, but also a definitive explanation for the inhibitory action of CsA and a new model of Rgg2 and Rgg3 regulatory interplay.

More specifically, in contrast to previous models describing Rgg2 as a transcriptional activator and Rgg3 as transcriptional repressor, we propose that Rgg2 and Rgg3 proteins are both transcriptional activators; however, in species encoding multiple Rgg proteins that are in competition for the same Rgg-box, an Rgg protein whose activation requires a significantly greater concentration of SHP can function as a repressor by sterically blocking access to the shared DNA binding site. To our knowledge this is a new model of transcriptional repression, where a DNA binding-competent yet *inactive* transcriptional activator functions as a repressor by inhibiting the access of another activator to a shared promoter.

## Results

### Cryo-EM Structure of the Rgg3_St_-SHP3 Complex and X-ray Crystal Structure of Rgg3_St_

The molecular weight of Rgg proteins (66 kDa dimer) approaches the lower limit of the high-resolution structures that have been solved using cryo-EM (14, 15). Despite this fact, when crystallization of SHP-bound Rgg proteins proved intractable we evaluated the use of single-particle cryo-EM for determination of the complex structure. We observed that Rgg3_St_ behavior was robust under cryogenic conditions, resulting in excellent particle distribution (Fig. S1A). Then, using the workflow described in Fig. S1B, we successfully determined the structure of Rgg3_St_-SHP3 to 3 Å resolution (Figs. 1 A-B, and S1; and Table S1). In addition to the cryo-EM structure of Rgg3_St_-SHP3, we also determined the X-ray crystal structure of Rgg3_St_ alone to 2.20 Å resolution (Fig. 1 C-D and Table S2). In both structures, the electron density corresponding to the DNA binding domain is uninterpretable, suggesting that the DNA binding domain is mobile, adopting numerous conformations relative to the repeat domain. In contrast, the electron density corresponding to the Rgg3 repeat domain and bound SHP3 are readily interpretable (Fig. S3 A-B). As detailed below, these structures reveal the contacts responsible for Rgg-SHP interaction specificity, the SHP-triggered Rgg repeat domain conformational change, the mechanistic basis underlying the phenotypes of numerous *rgg* mutations (12), and the mechanism of action of CsA, a previously identified inhibitor of SHP-triggered Rgg function (13, 16).

**Fig 1.**
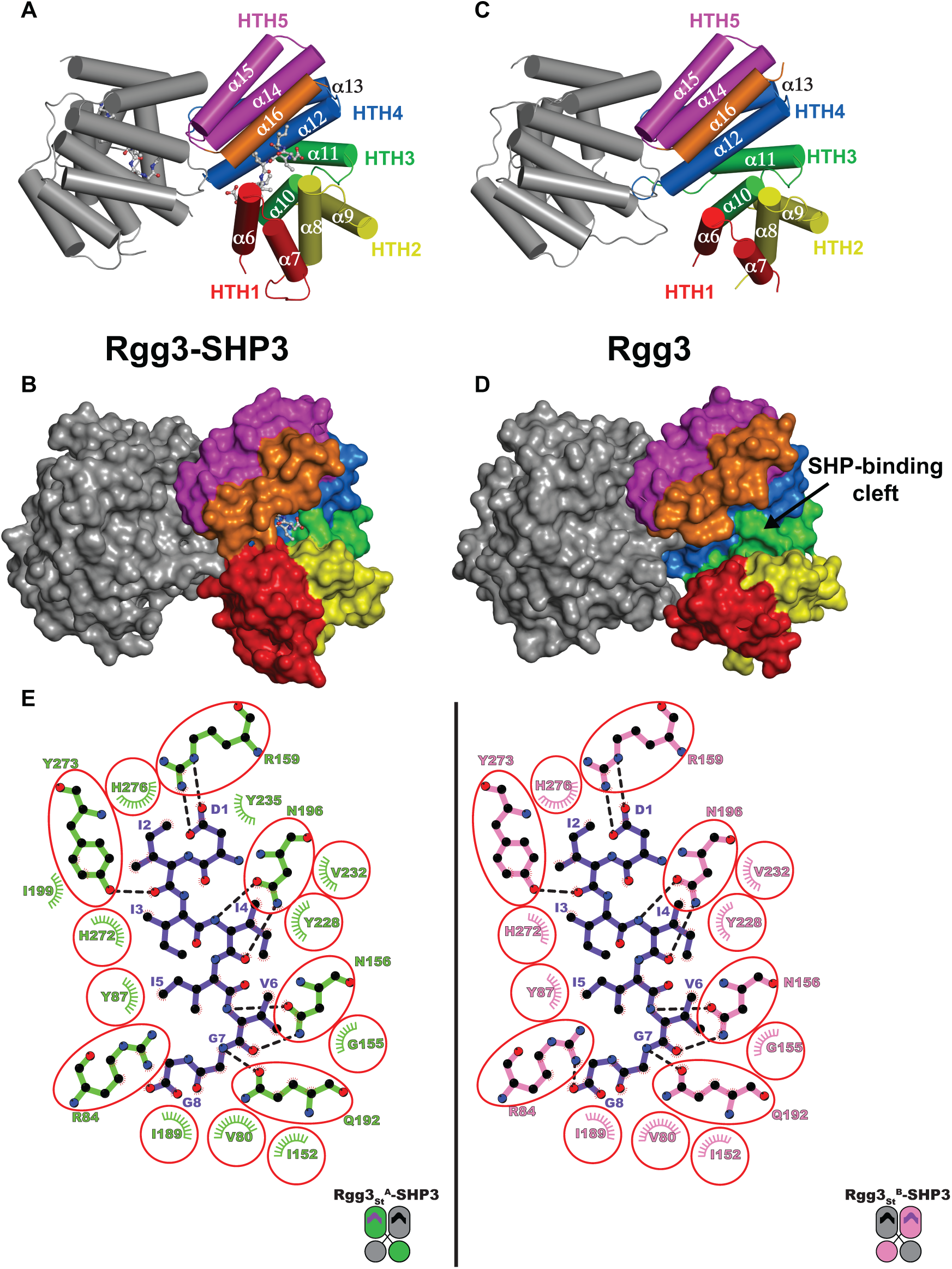
Cryo-EM structure of Rgg3_St_-SHP3 and X-ray crystal structure of Rgg3_St._ (A) The cryo-EM structure of Rgg3_St_ (rainbow-colored cylinders depict the repeat domain α-helices of one protomer while the other protomer is colored grey) complexed with SHP3 (ball and stick models). (B) The cryo-EM structure of Rgg3_St_ (rainbow surface) bound to SHP3 (ball and stick model). (C) The X-ray crystal structure of Rgg3_St_ alone. (D) The X-ray crystal structure of Rgg3_St_ alone (rainbow surface). Pheromone binding triggers Rgg3_St_ SHP-binding cleft closure. (E) Schematic representation of SHP3 interactions with Rgg3^A^_St_ (colored green, left) and Rgg3^B^_St_ (colored magenta, right). SHP3 is depicted as purple bonds, hydrogen bonds are depicted as black dashed lines, hydrophobic contacts mediated by Rgg3^A^_St_ and SHP3 are shown as green semicircles with radiating lines, while hydrophobic contacts mediated by Rgg3^B^_St_ and SHP3 are shown as pink semicircles with radiating lines. Red circles indicate SHP3-binding residues in common to Rgg3^A^_St_ and Rgg3^B^_St_. The schematic was produced with LigPlot+ (20). HTH, helix-turn-helix.

*S. thermophilus* SHP3 (amino acid sequence DIIIIVGG) is an archetype SHP peptide, containing an N-terminal acidic residue followed by seven hydrophobic amino acids (Fig. 1 A-B, and E). The Rgg3_St_-SHP3 structure shows that SHP3 binds to the concave surface of the right-handed superhelical repeat domain (Fig. 1 A and E). Structural comparison of Rgg3_St_-SHP3 and Rgg3_St_ alone shows that upon binding SHP3, the repeat domain compresses along its superhelical axis, transitioning from an open to closed conformation, largely burying SHP3 in the repeat domain core and generating new contacts between previously distant repeat domain helices, e.g., α6 and α16 (Fig. 1 A-D).

The Rgg3_St_-SHP3 interactions are extensive (Fig. 1E). The SHP3 N-terminal aspartate forms a salt-bridge with the side chain of Rgg3_St_ Arg159, which is highly conserved in Rgg2 and Rgg3 proteins associated with group I SHPs (i.e., SHPs containing an N-terminal aspartate)(Figs. 1E and S4)(10). While the side chains of the remaining SHP3 residues mediate hydrophobic interactions, numerous SHP3 main chain nitrogen and carbonyl atoms mediate H-bonds with Rgg3_St_ side chains. Furthermore, the SHP3 carboxyl-terminus forms a salt bridge with Rgg3_St_ residue R84, which is highly conserved in Rgg2 and Rgg3 proteins associated with both group I and II SHPs (Figs. 1E and S4)(10).

Consistent with the observed Rgg3_St_-SHP3 interactions, we previously found that alanine substitutions in Rgg2_Sd_ residues R81, R150, R153, and N190 equivalent to Rgg3_St_ residues, R84, R156, R159, and N196 caused loss of SHP-trigged function in *S. pyogenes* (13). Together, these data show that SHP3 is anchored to Rgg3_St_ via salt bridges at the SHP3 N- and C-terminus; and that interactions throughout SHP3 bind it to Rgg3_St_ via hydrophobic contacts mediated by SHP3 side chain atoms and H-bonds mediated by SHP3 main chain atoms. These extensive interactions explain how the relatively short SHPs bind with high affinity and specificity to their cognate Rgg proteins (e.g., *S. pyogenes* Rgg3-SHP3 *K*_d_ = 1.88 μM) (8).

### X-ray Crystal Structure of the Rgg2_Sd_-Rgg Box Complex

To determine how Rgg proteins recognize target DNA promoters, we first determined the X-ray crystal structure of Rgg2_Sd_ in complex with Rgg-box DNA (Fig. 2). The 2.8 Å resolution structure was determined using phases obtained by molecular replacement (see Experimental Procedures and Table S2). The crystallographic asymmetric unit contains an Rgg2_Sd_ domain-swapped dimer and one molecule of double-stranded DNA (dsDNA). We refer to the Rgg2_Sd_ protomers as Rgg2^A^_Sd_ and Rgg2^B^_Sd_ (Fig. 2 A and B); and, as necessary, we indicate whether nucleotides belong to the top or bottom strands of the Rgg box by the absence or presence of the prime symbol, e.g., C^9^ and G^19^′, respectively (Fig. 2 E).

**Fig 2.**
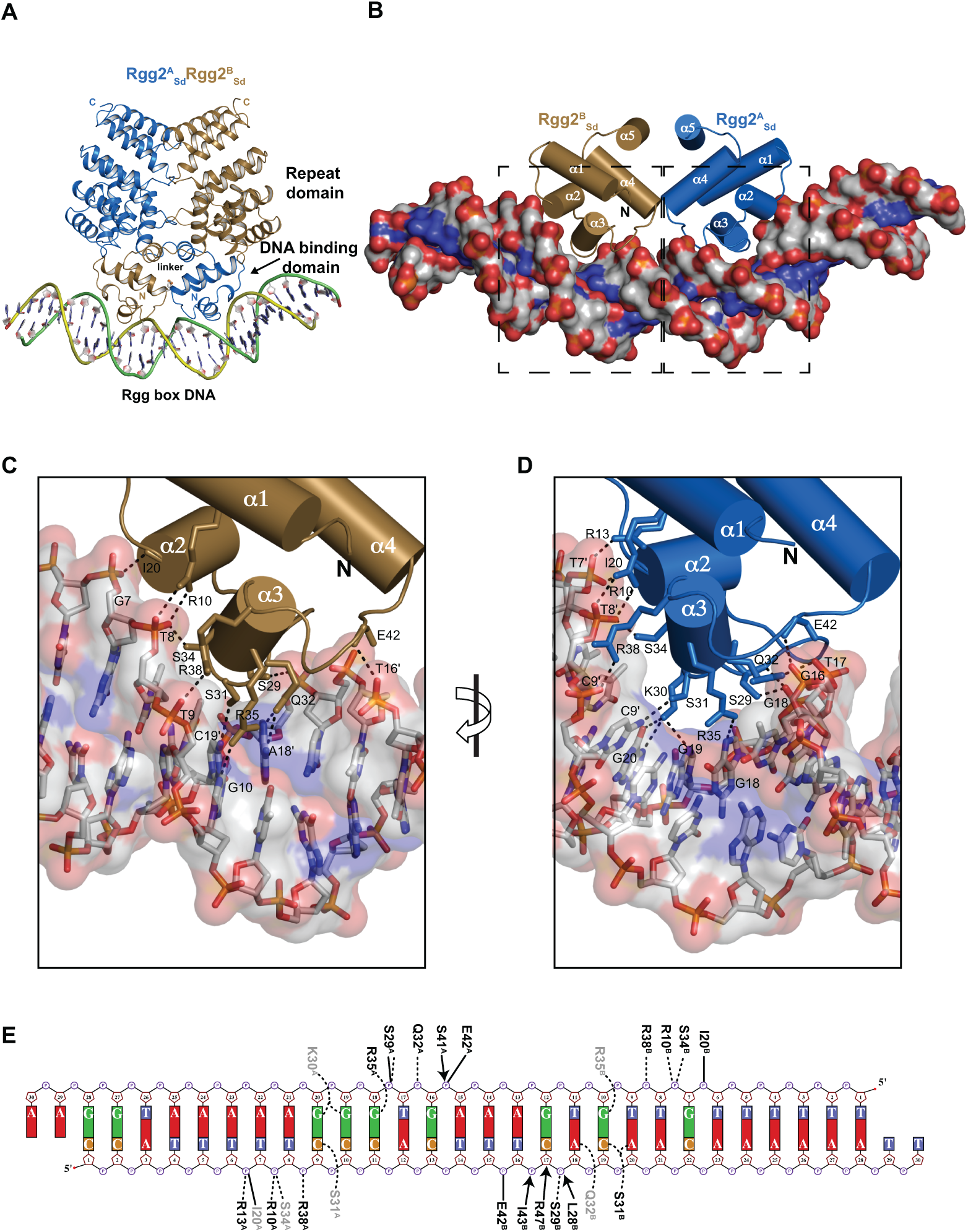
Crystal structure of Rgg2_Sd_ bound to Rgg2-box DNA. (A) Rgg2_Sd_ dimer bound to Rgg2-box DNA. The intermolecular disulfide bond connecting the DNA binding domains is depicted as balls and sticks. (B) Isolated view of the Rgg2_Sd_ DNA binding domains (cylindrical helices) in complex with Rgg-box DNA (surface rendering colored by element). (C and D) Expanded view of the areas enclosed by the dashed black rectangles on the left or right in panel B, respectively. The structures were rotated slightly to provide a clear view of the Rgg2_Sd_-Rgg box interactions. Rgg-box is depicted as a semi-transparent surface and sticks. Rgg2_Sd_ side chains or main chains that interact with Rgg-box are labeled and shown as sticks. Hydrogen bonds are depicted as dashed lines. (E) Rgg2_Sd_-Rgg box interaction schematic. Solid and dashed lines indicate H-bond interactions between Rgg box and the Rgg2_Sd_ main chain and side chains, respectively. Arrows represent intermolecular hydrophobic interactions (<3.35 Å). Sidechains with ambiguous electron density are shaded grey. DNA phosphate is depicted as circles, ribose sugars as pentagons, and nucleotide bases as rectangles. Portions of the schematic were generated using NUCPLOT (21).

Structural alignment of the Rgg2_Sd_ protomers from the Rgg2_Sd_-Rgg box structure shows that the proteins are in similar conformations (rmsd for modeled C_α_ = 0.98 Å) and related by a two-fold symmetry axis; however, it is important to note that the Rgg2_Sd_-Rgg box structure is intrinsically asymmetric due to the fact that the promoter DNA is non-palindromic (Fig. 2E). Comparison of the Rgg2_Sd_-Rgg box and Rgg2_Sd_ (PDB ID 4YV6) structures shows that Rgg2_Sd_ is dimeric in the presence or absence of DNA, and that Rgg2_Sd_ adopts nearly identical conformations in both structures (rmsd for modeled C_α_ = 0.50 - 0.60 Å for measured pair-wise comparisons of Rgg2_Sd_ dimers). Consistent with their structural similarly, the intermolecular disulphide bond we previously identified in the structure of Rgg2_Sd_ alone is also present in the crystal structure of Rgg2_Sd_-Rgg box as well as in solution (Figs. 2A and S5)(13).

The DNA binding domains of both Rgg2^A^_Sd_ and Rgg2^B^_Sd_ interact with the Rgg box, which is B-form and bent 47.8° (Fig. 2A)(17). The Rgg2^A^_Sd_-DNA and Rgg2^B^_Sd_-DNA interfaces bury a similar amount of surface area 1,210.0 Å^2^ and 1,243.7 Å^2^, respectively. While non-specific interactions with the Rgg-box sugar-phosphate backbone are mediated by amino acids distributed throughout much of the DNA binding domains including helices α1-α4 and their intervening loops (Fig. 2B-E), only residues in the DNA binding domain helix α3 mediate base-specific contacts (Fig. 2B-E). Nucleotide base-specific hydrogen bonds are mediated by the side chains of Rgg2^A^_Sd_ residues K30 with G^19^′ and G^20^′; S31 with C^9^; and R35 with G^18^′; as well as Rgg2^B^_Sd_ residues S31 with C^19^ and A^20^; Q32 with A18; and R35 with G10.

### X-ray Crystal Structures of Rgg3_St_-Rgg Box and Rgg3_St_

In the existing model of Rgg function, Rgg2 and Rgg3 proteins are transcriptional activators and repressors, respectively. We proposed that structural comparison of Rgg2 and Rgg3 proteins in complex with DNA would reveal the mechanistic basis of this functional difference. Thus, with the structure of Rgg2_Sd_-Rgg box in hand, we determined the X-ray crystal structure of *S. thermophilus* Rgg3 (Rgg3_St_) in complex with Rgg-box DNA to 3.20 Å resolution (Fig. S6 and Table S2).

Overall, the structure of Rgg3_St_-Rgg box (Fig. S6) is very similar to the structure of Rgg2_Sd_-Rgg box (Fig. 2) with an rmsd for modeled C_α_ = 1.11 Å. Consistent with the striking amino acid conservation of their α3 helices (Fig. S4), Rgg3_St_ employs many of the same residues as Rgg2_Sd_ to bind DNA specifically and non-specifically (Figs. S4 and S6). However, consistent with the lower resolution of the Rgg3_St_-Rgg box structure, the quality of the electron density corresponding to its Rgg box DNA and interacting Rgg3_St_ side chains was comparatively lower than that of the Rgg2_Sd_-Rgg box structure (Fig. S7), thus, side chain rotameric configurations were, in some cases, less definitive for Rgg3_St-_Rgg box than for Rgg2_Sd_-Rgg box. For example, Rgg3_St_ residue R10 likely forms bidentate H-bonds to T7′ (equivalent to Rgg2_Sd_-box DNA T8′); however, as modeled, to form these bonds the Rgg3_St_ R10 side chain would be required to shift approximately 0.6 Å (Figs. 2E, S6, and S7). In sum, Rgg2_Sd_ and Rgg3_St_ share a striking overall structural similarity that is consistent with the revised model of Rgg2/3 function presented below.

### Cyclosporin A and SHP Bind to Overlapping Sites on Rgg Proteins, and Cyclosporin A Sterically Blocks the SHP-Induced Rgg Conformational Change

We previously demonstrated that CsA is an inhibitor of SHP binding to Rgg proteins and that Rgg binding to DNA is unaffected by the presence of CsA (13). We also determined the X-ray crystal structure of CsA in complex with Rgg2_Sd_, which showed that CsA can bind in two different conformations to the repeat domain core (Figs. 3A and S8), and we found in genetic studies that many of the Rgg2_Sd_ residues that interact with CsA are also required for SHP2 to activate Rgg2_Sd_. Thus, we proposed that the CsA- and SHP-binding sites might overlap. Based on a structural comparison of Rgg3_St_-SHP3 (Fig. 3B) and Rgg2_Sd_-CsA (Figs. 3C and S8), we conclude that CsA and SHP compete to bind overlapping sites on the concave surface of the Rgg repeat domain core.

**Fig 3.**
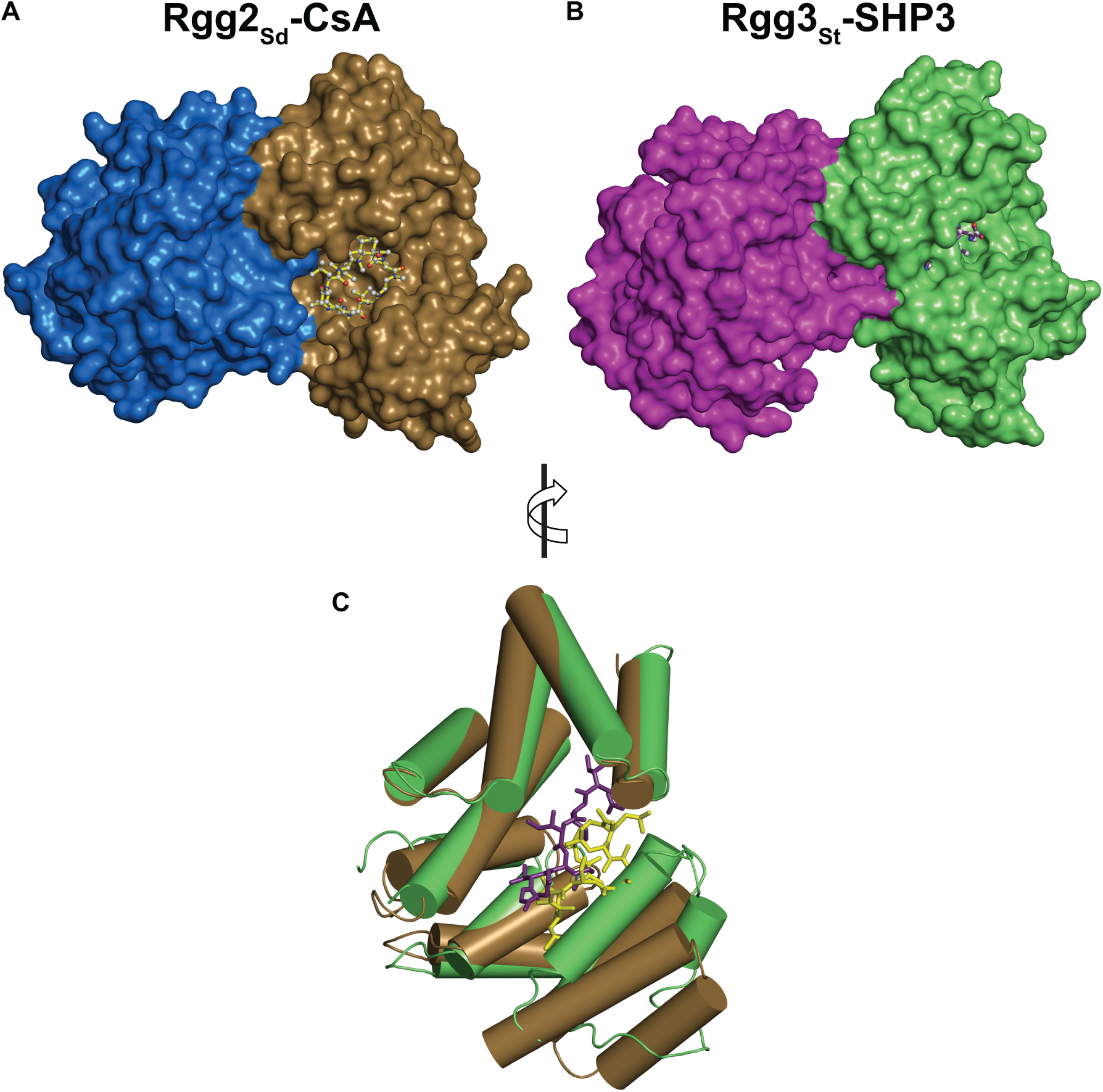
SHP3 and cyclosporin A bind to overlapping sites on their target Rgg proteins. (A) X-ray crystal structure of Rgg2_Sd_ dimer complexed with CsA (ball and stick model) (PDB 4YV9). (B) Cryo-EM structure of Rgg3_St_ complexed with SHP3 (ball and stick model). For clarity, the Rgg2_Sd_ DNA binding domains are not shown. (C) Structural alignment of Rgg2^B^_Sd_ (brown cylinders) bound to CsA^F^ (yellow sticks) (PDB 4YV9) and Rgg3^A^_St_ (green cylinders) bound to SHP3 (purple sticks). For clarity, only one monomer of each Rgg2/3 dimer is shown, and the Rgg2^B^_Sd_ DNA binding domain is omitted.

Interestingly, Rgg2_Sd_-CsA, Rgg2_Sd_-Rgg box, and Rgg3_St_-Rgg box are in essentially identical conformations, i.e., in each case Rgg2_Sd_ or Rgg3_St_ is in the repeat domain open, i.e., SHP-accessible (if not for the presence of CsA), and DNA binding proficient conformation. Based on the alignment of Rgg3_St_-SHP3 and Rgg2_Sd_ bound to CsA (Figs. 3C and S8), it is now clear how CsA prevents Rgg proteins from adopting the closed (transcriptionally active) conformation. CsA not only competes with SHP for access to their overlapping binding site in the repeat domain core, CsA also acts as a wedge, locking the Rgg repeat domain in the open (transcriptionally inactive) conformation.

### Rgg2 and Rgg3 Proteins are Transcriptional Activators

In the course of Rgg2 and Rgg3 functional studies, we routinely employ a luciferase reporter transcriptional bioassay (18). In this bioassay, the luciferase genes are under the control of the Rgg2- and Rgg3-regulated promoter P*shp3*. Using these bioassays, we previously demonstrated that Rgg2_Sp_ is an activator, triggered by SHPs. We also showed that Rgg3_Sp_ is a transcriptional repressor, de-repressed by SHP peptide. However, we show here that at concentrations of SHP3 higher than previously examined Rgg3_Sp_ is in fact a comparatively weak activator at P*shp3* (Fig. 4 A-B).

**Fig 4.**
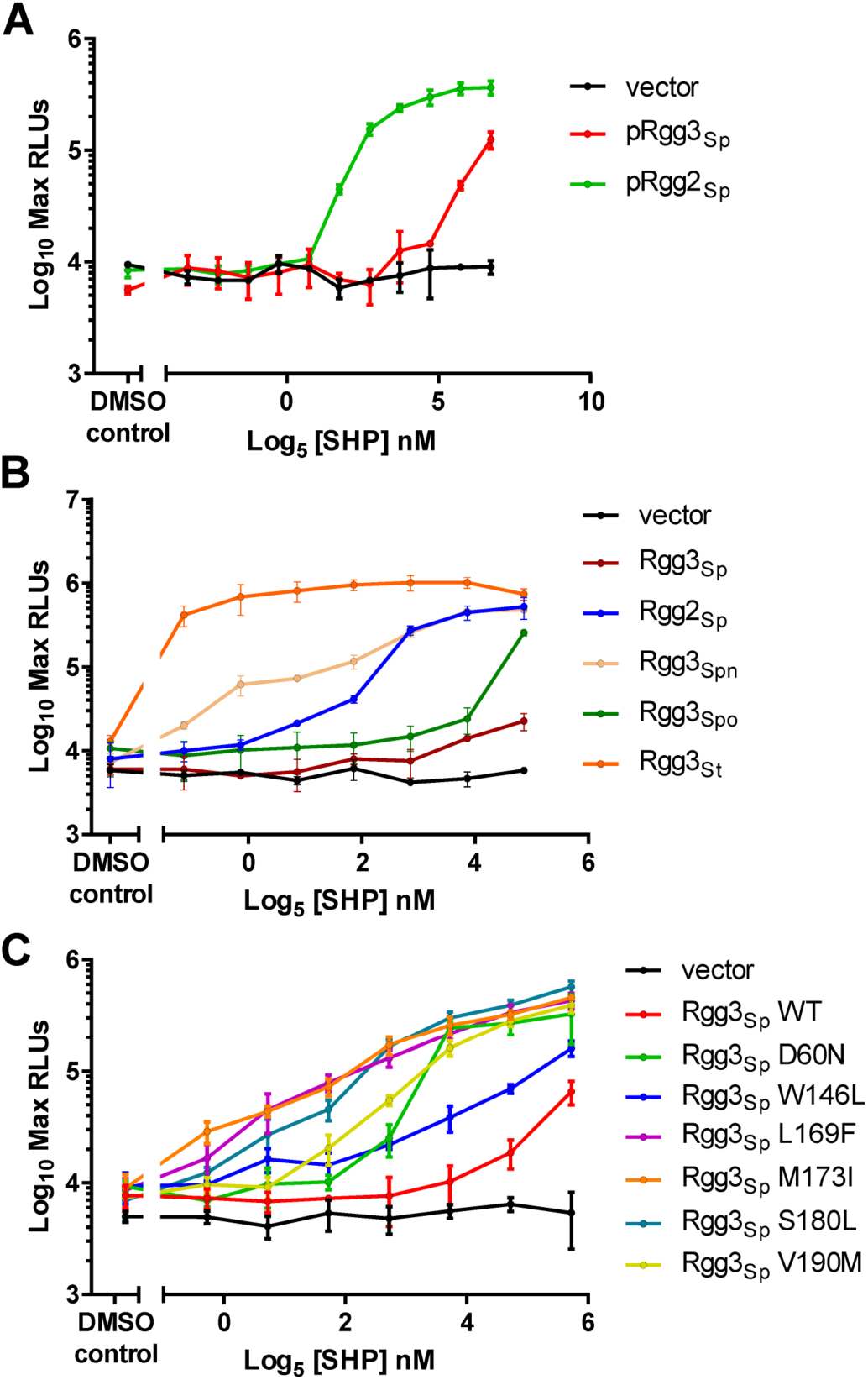
Rgg2 and Rgg3 proteins are transcriptional activators. Luciferase luminescence assays reporting chromosomally inserted *shp3* promoter fused to *luxAB* in a GAS host null for *rgg2, rgg3, shp2, shp3* (strain KMT101). (A) Rgg2_Sp_ and Rgg3_Sp_ were individual expressed from plasmid pLZ12-spec (vector) in KMT101 and luciferase reporter activity was measured in response to SHP dosage. (B) Rgg homologs from *S. pyogenes* NZ131 (Sp), *S. pneumoniae* str. R6 (Spn), *S. porcinus* str. Jelinkova 176 (Spo), and *S. thermophilus* CNRZ1066 (St) were expressed in KMT101 as above. (C) Rgg3_Sp_ mutants expressed from plasmid pLZ12-spec (vector) in KMT101. Error bars indicate standard deviation from three independent biological replicates.

When we carried out similar analysis using Rgg3_St_, we discovered that it is a remarkably strong activator at P*shp3* even at low concentrations of SHP3 (Fig. 4 B). Motivated by the finding that Rgg3_Sp_ and Rgg3_St_ are transcriptional activators (when, again, in previous models they were suspected to be repressors), we examined additional Rgg3 proteins, namely *S. pneumoniae* Rgg3 (Rgg3_Spn_) and *S. porcinus* Rgg3 (Rgg3_Spo_). Both proteins activated transcription at P*shp3* (Fig. 4 B). Thus, all Rgg2 and Rgg3 proteins examined to date are SHP2/3 responsive transcriptional activators. How Rgg3_Sp_ also functions as a transcriptional repressor in the context of Rgg2_sp_ is discussed below.

In genetic selections to generate Rgg3_Sp_ mutations that enhance its transcriptional activity, we identified Rgg3_Sp_-M173I (Fig. 4C). We previously found that the analogous substitution in Rgg2_Sp_ (M167I) caused SHP-independent Rgg2_Sp_ activation (12). Given the similar effect of these mutations in Rgg2_sp_ and Rgg3_Sp_, additional residues we previously found to confer constitutive Rgg2_Sp_ activity were constructed at corresponding positions in Rgg3_Sp,_ Indeed, these mutations significantly increased Rgg3_Sp_ transcriptional activity *in vivo* (Fig. 4C). None of the wild-type residues substituted in the gain-of-function mutants interact directly with SHP3, and we demonstrated previously for Rgg2_Sp_ that they do not increase SHP- or DNA-binding affinity (12). Thus, the mutants appear to function allosterically, likely stabilizing Rgg2 and Rgg3 proteins in an activated (closed) conformation. Together, the above results are consistent with a model where Rgg2 and Rgg3 proteins are activators that function in a mechanistically similar if not identical manner.

### *In Vitro* and *In Vivo* Functional Analysis of Conserved Rgg2-Rgg box and Rgg3-Rgg box interactions

With an Rgg3_St_ luciferase reporter transcriptional bioassay in hand (Fig. 4), we sought to confirm the mechanistic and biological importance of the protein-DNA contacts observed in both the Rgg2_Sd_-Rgg box and Rgg3_St_-Rgg box crystal structures. In addition to *in vivo* genetic Rgg3_St_ P*shp3* luciferase reporter transcriptional bioassays (Fig. 5A), we carried out *in vitro* electrophoretic mobility shift assays (EMSAs) (Fig. 5B). More specifically, for these *in vivo* and *in vitro* loss-of-function studies we mutated to alanine selected residues in Rgg3_St_ that are conserved in Rgg2 and Rgg3 proteins and observed to contact DNA in both the Rgg2_Sd_-Rgg box and Rgg3_St_-Rgg box crystal structures (Figs. 2E and S6E).

**Fig 5.**
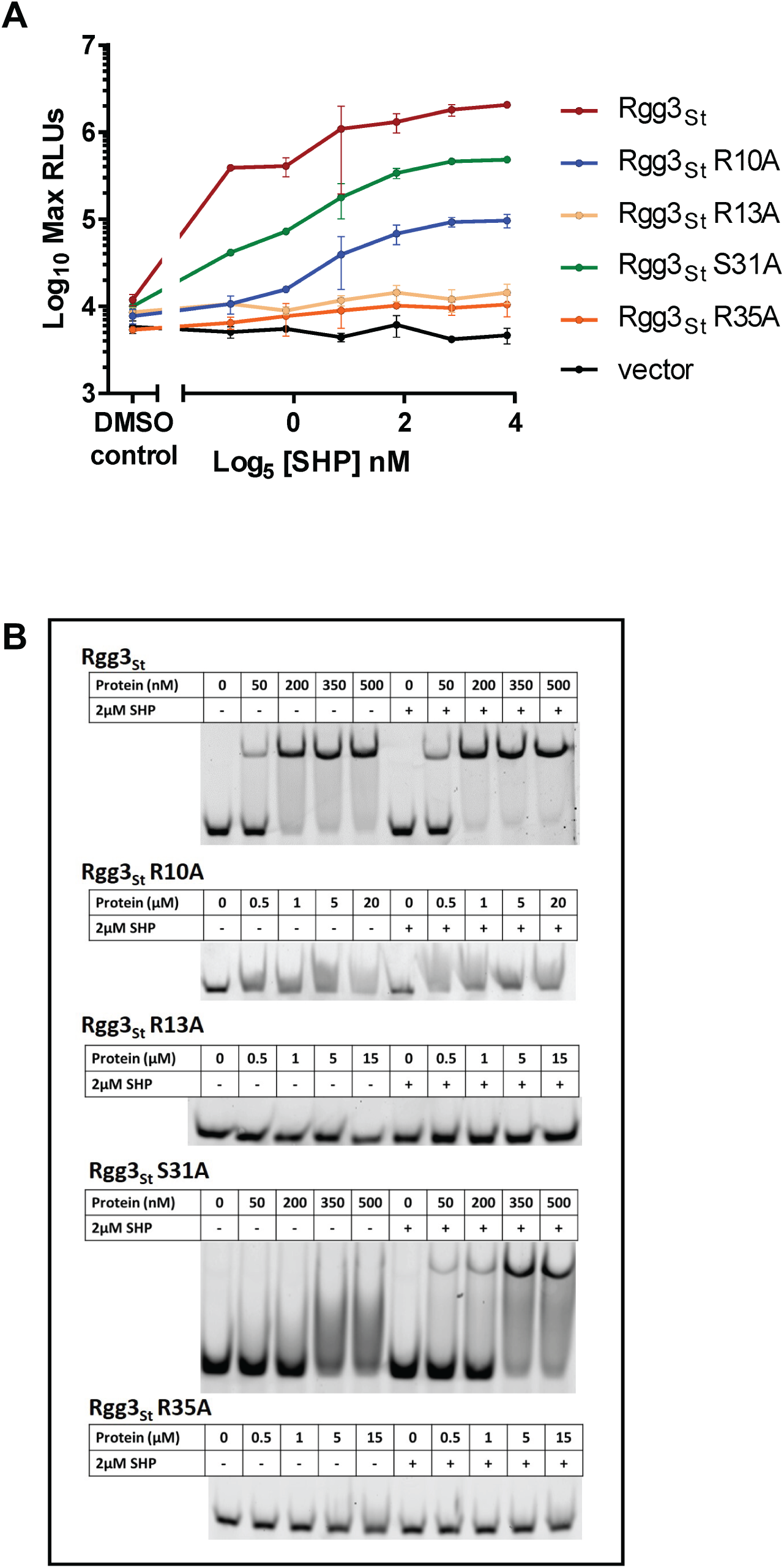
Functional importance of conserved DNA-binding Rgg3_St_ amino acids. Comparison of Rgg3_St_ alanine-substitution mutants to wild-type Rgg3_St_ using an *in vivo* Pshp3 luciferase reporter assay (A) and *in vitro* using EMSAs (B). The mutations were introduced in a *S. pyogenes* strain containing *rgg3*_St_ cloned from a gene synthesized codon-optimized for expression in *E. coli*. The EMSAs shown in (B) are representative of 2-3 independent experiments.

In sum, in comparison to wild-type Rgg3_St_, the Rgg3_St_-R13A and Rgg3_St_-R35A mutants displayed a complete loss of function (Fig. 5A), i.e., they showed no response whatsoever to SHP3. Rgg3_St_-S31A and Rgg3_St_-R10A, however, displayed intermediate phenotypes, i.e., they had SHP3-dependent activity but were far less active than wild-type Rgg3_St_. Consistent with these results, Rgg3_St_-R13A and Rgg3_St_-R35A did not bind to the labeled DNA probe in the EMSA, while Rgg3_St_-R10A and Rgg3_St_-S31A displayed severe DNA-binding defects (Fig. 5B). Interestingly, the Rgg3_St_-S31A defect was partially compensated for by the presence of SHP3 *in vitro*, which may also explain why its phenotype was less severe than that of Rgg3_St_-R10A *in vivo* (Fig. 5 A-B).

## Discussion

The archetype RRNPP family member Rgg proteins have emerged as one of the most important regulatory transcription factors in streptococci, controlling virulence, antimicrobial resistance, and biofilm formation (2-5). We propose a revised, unifying model of Rgg2/Rgg3 function based on a combination of biochemical, genetic, X-ray crystallographic, and cryo-EM studies (Fig. 6). While there may be Rgg proteins that prove to be exceptions to this model, we propose that both Rgg2 and Rgg3 proteins are in fact transcriptional activators, and Rgg3 proteins are not inherently transcriptional repressors as previously described. In bacteria such as Group A Streptococcus that have two or more Rgg proteins in competition for binding to identical promoter elements, the Rgg proteins can also function as repressors sterically blocking access to Rgg box DNA if at least one of these receptors has relatively higher activation requirements. For example, if their *in vivo* activation threshold requires different SHP concentrations, an inactive Rgg could inhibit the DNA binding of other activated Rggs. To our knowledge, a similar gene expression regulatory scenario has not been previously described but in all likelihood is not unique to streptococci.

**Fig. 6.**
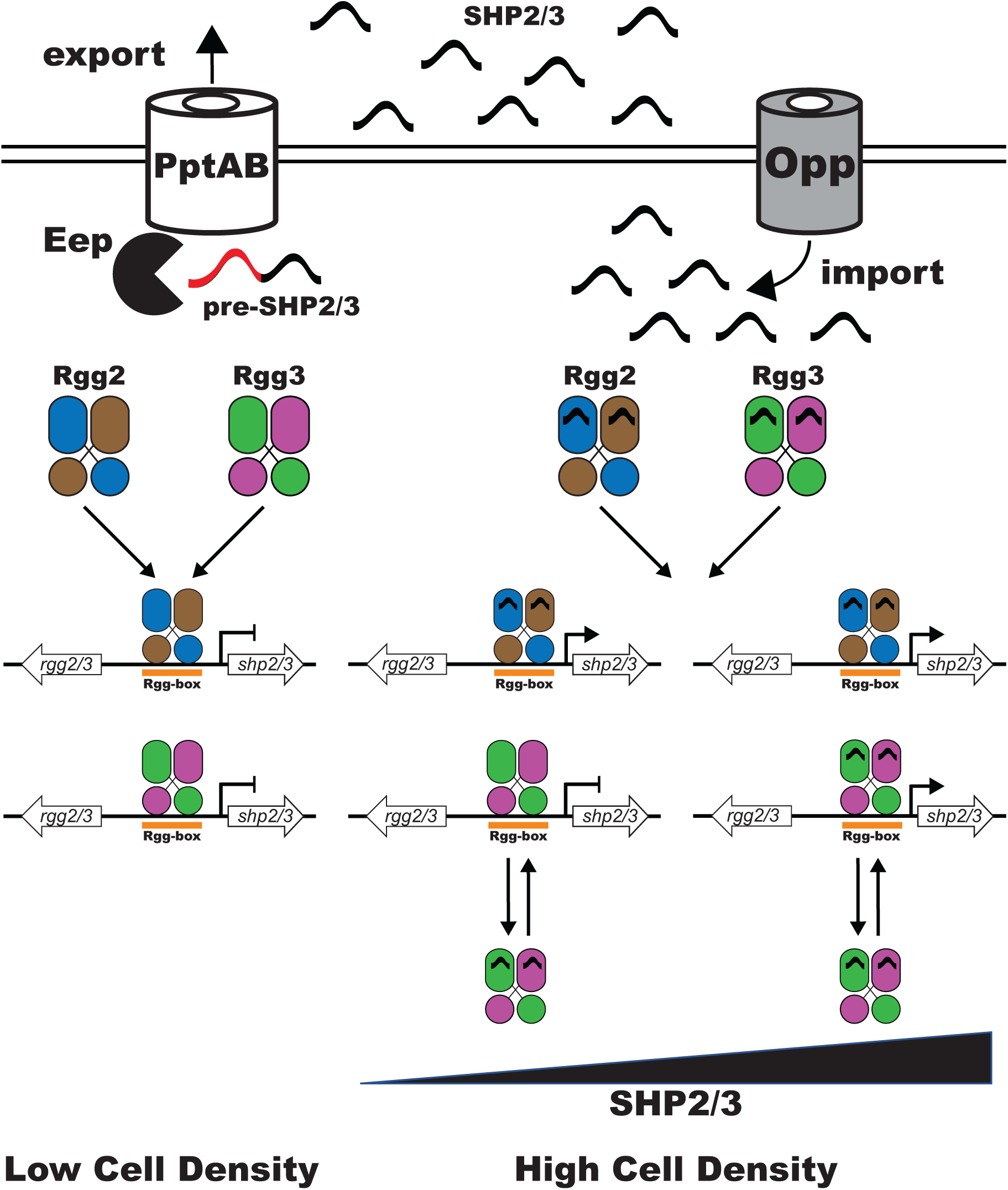
Updated model of Rgg-SHP quorum-sensing signaling. In contrast to the existing model of Rgg-SHP signaling where Rgg2 proteins are transcriptional activators and Rgg3 proteins are transcriptional repressors, in the updated model Rgg2 and Rgg3 proteins are both SHP-dependent transcriptional activators. However, in streptococci such as *S. pyogenes* that contain more than one Rgg-SHP signaling system, crosstalk, competition, and differential activation requirements result in complex interplay. For example, Rgg proteins such as Rgg3_Sp_ can function as repressors if they are in competition for promoter access with another Rgg protein such as Rgg2_sp_ whose activation requires comparatively less SHP. Furthermore, in addition to triggering its transcriptional activity, SHP2 or SHP3 binding to Rgg3_Sp_ also appears to moderately reduce its affinity for DNA.

Furthermore, we have identified one Rgg, Rgg3_Sp_, whose DNA binding affinity is moderately decreased upon binding SHP2 or SHP3 (19). While Rgg3_Sp_ is a SHP2/3-dependent transcriptional activator, consistent with the fact that it functions primarily as a transcriptional repressor competing with Rgg2_Sp_ for binding SHP2/3 and Rgg box DNA, Rgg3_Sp_ transcriptional activation requires higher concentrations of SHP2/3 than does Rgg2_sp_ (Fig. 4). We propose that in the presence of what appears to be a largely redundant Rgg, Rgg2_Sp,_ there was little pressure to maintain Rgg3_Sp_ SHP-driven activator function. Thus, at physiological levels of SHP2/3, Rgg3_Sp_ serves as a repressor, tuning the cellular threshold for SHP2/3 response.

What is the molecular basis of SHP-Rgg binding specificity? Based on the Rgg3_St_-SHP3 cryo-EM structure, the conserved Group I SHP N-terminus Asp, the conservation of the Rgg position equivalent to Rgg3_St_ amino acid 159 in Rgg proteins associated with Group I SHPs as Arg, and the fact that all SHPs likely have a C-terminal carboxyl group, we propose the following model of Group I SHP molecular recognition. Group I SHPs recognize their cognate Rgg proteins by scanning their surface for a largely hydrophobic complementary shape. Following this initial recognition event whose specificity is determined primarily by the SHP side chains, the SHP-Rgg interaction is anchored by H-bonds between the SHP main chain and Rgg side chains as well as critical hotspot interactions. The hotspot interactions are salt bridges at the SHP N- and C-termini. The N-terminal SHP salt bridge is mediated by its conserved Asp1 side chain and a conserved Arg side chain in its target Rgg (at the position equivalent to Rgg3_St_ R159)(Fig. 1E). We previously found that a mutation here eliminated SHP-triggered activity in Rgg2_Sp_ (12, 13). The C-terminal SHP salt bridge is mediated by the C-terminal carboxy group and a conserved Arg side chain in its cognate Rgg (at the position equivalent to Rgg3_St_ 84). We similarly found that a mutation here eliminated SHP-trigged Rgg2_Sp_ activity (13). We suspect that SHPs containing Glu at their N-terminus, e.g., Group II SHPs, recognize their cognate Rgg proteins using a similar mechanism; however, their binding mode in the Rgg repeat domain core is going to be somewhat different because there is no basic residue in their associated Rgg proteins at the position equivalent to Rgg3_St_ R159.

SHP3 binding to Rgg3_St_ induced a large conformational change, compressing the repeat domain along its helical axis and closing the otherwise open repeat domain cleft. How does this SHP-triggered conformational change regulate Rgg transcriptional activation? Our X-ray crystallographic and cryo-EM structures of Rgg proteins point to the fact that the repeat domain and DNA binding domain are connected by a flexible linker and that the repeat domain and DNA binding domain move relative to one another. While the structure of an Rgg in complex with DNA and SHP will be required to answer this question, we propose that in response to SHP binding, the closed repeat domain dimer rotates relative to the DNA binding domain, which undergoes little conformational change in part due to the fact that the DNA binding domains are linked by intermolecular disulphide bonds. The SHP-bound, rotated repeat domain exposes a surface to RNA polymerase, activating transcription.

As a result of these structure-function studies, we are now well positioned to design pharmacological inhibitors of Rgg function. In addition to designing new inhibitors based on SHP mimetics, our results make clear how the cyclic undecapeptide CsA functions to inhibit Rgg proteins. Based on our studies, the stage is set for the synthesis of non-immunosuppressive CsA derivatives that leverage critical interactions observed in the Rgg3_St_-SHP and Rgg2_Sd_-CsA structures.

## Materials and Methods

The Rgg3_St_-SHP3 structure was determined using single-particle cryo-electron microscopy (cryo-EM). The structures of Rgg3 alone, Rgg2_Sd_-Rgg box, and Rgg3_St_-Rgg box were determined using X-ray crystallography.

Details of the methods are presented in *SI Appendix, SI Materials and Methods*.

## Acknowledgments

We thank Emre Firlar for assistance collecting cryo-EM data, Ned Wingreen for helpful discussions, and Atul Khataokar for critical review of the manuscript. X-ray diffraction data were collected at the Stanford Synchrotron Radiation Light Source. Use of the Stanford Synchrotron Radiation Lightsource, SLAC National Accelerator Laboratory, is supported by the U.S. Department of Energy, Office of Science, Office of Basic Energy Sciences under Contract No. DE-AC02-76SF00515. The SSRL Structural Molecular Biology Program is supported by the DOE Office of Biological and Environmental Research, and by the National Institutes of Health, National Institute of General Medical Sciences (P41GM103393). The contents of this publication are solely the responsibility of the authors and do not necessarily represent the official views of NIGMS or NIH. Support for this work was provided by National Institutes of Health Grants R01 AI125452 (M.B.N. and M.J.F.); by National Institutes of Health Grants R01 AI091779 (M.J.F.); by the Burroughs Wellcome Fund Investigators of Pathogenesis of Infectious Diseases (M.J.F.); and by the Chicago Biomedical Consortium with support from the Searle Funds at the Chicago Community Trust (M.J.F.).

## Data Deposition

Atomic coordinates and structure factors for Rgg3_St_-Rgg box, Rgg2_Sd_-Rgg box, and Rgg3 alone have been deposited in the Protein Data Bank under accession codes 61WF, 61WA, 61WE, respectively. Consistent with journal requirements, the atomic coordinates and cryo-EM map for Rgg3_St_-SHP3 will be deposited in the Protein Data Bank and in the Electron Microscopy Data Bank, respectively.

## Supplementary Information for

## Supplementary Materials and Methods

### Rgg2_Sd_ Production

*S. dysgalactiae rgg2* was cloned in pTB146 as previously described (1). His-Sumo-Rgg2_Sd_ was overexpressed in *E. coli* strain BL21(DE3) by first growing the cells at 37 °C in LB medium containing 100 μg/ml ampicillin to OD_600_ = 0.5 and then inducing expression with 0.5 mM isopropyl β-D-1-thiogalactopyranoside (IPTG) for 4 h at 25 °C. The cells were collected by centrifugation and lysed in Buffer A (50 mM Tris·HCl [pH 8.0], 500 mM NaCl, 10% glycerol) supplemented with 20 μg/ml DNase. Lysate supernatant was applied to His-60 Ni resin (Clontech) equilibrated in Buffer A. His-Sumo-Rgg2_Sd_ was eluted by washing the column with increasing amounts of imidazole and analyzed for purity using SDS-PAGE. Eluted protein was combined with 1.25 mg of the SUMO protease Ulp1 and dialyzed against 2 L of Buffer B (20 mM sodium phosphate buffer [pH 8.0], 150 mM NaCl, 10 mM β-ME, and 0.1% Triton X-100) overnight at 4 °C. The next day the protein was passed over His-60 Ni resin to separate the cleaved tag from the Rgg2_Sd_ protein. The resulting cleaved protein, Rgg2_Sd_, was pooled, concentrated by ultrafiltration through a 10 kDa filter, and further purified by gel filtration using a Superdex 200 (GE Healthcare) 16/70 column equilibrated in Buffer C (20 mM Tris·HCl [pH 8.0], 150 mM NaCl). Rgg2_Sd_ was concentrated using a 10 kDa MWCO centrifugal filter device and stored at −80 °C.

### Rgg3_St_ Production

*S. thermophilus rgg3* (Rgg3_St_) was synthesized optimized for expression in *E. coli* (Genscript, Inc.). The Rgg3_St_ expression construct encodes the 284 amino acids from GenPept entry WP_011225998.1. A PCR product was generated using Phusion High-Fidelity DNA Polymerase (New England Biolabs) and the primer pair His-Sumo-Stherm_F and His-Sumo-Stherm_R (Table S3). This PCR product was cloned into the SapI and XhoI sites of pTB146 using the Gibson Assembly method (New England Biolabs) to give His-Sumo-Rgg3_St_. Growth, overexpression, and purification followed the same protocol as used to produce Rgg2_Sd_ with the addition of 5 mM DTT to Buffers A and C. Rgg3_St_ was concentrated using a 10 kDa MWCO centrifugal filter device and stored at −80 °C.

### Rgg2_Sd_ and Rgg3_St_ Mutation and Production

Rgg2_Sd_ mutant C45S was generated *de novo* in His-Sumo-Rgg2_Sd_ using the QuickChange Lightning site-directed mutagenesis kit (Agilent) and verified by DNA sequencing. Rgg3_St_ mutants R10A, R13A, S31A, R35A, and C45S were regenerated *de novo* in His-Sumo-Rgg3_St_ using the QuickChange Lightning site-directed mutagenesis kit (Agilent) and verified by DNA sequencing. Growth, overexpression, and purification followed the same protocols described for Rgg2_Sd_ and Rgg3_St_.

### Rgg3_St_-SHP3 Cryo-EM Structure Determination

Rgg3_St_ was prepared as described above but during the final gel filtration step using a Superdex 200 (GE Healthcare) 16/70 column, the equilibration buffer contained 150 mM NaCl and 20 mM HEPES [pH 8.0]. SHP3 (DIIIIVGG) was obtained from LifeTein, LLC and resuspended in DMSO. Rgg3_St_-SHP3 complex was prepared by incubating 30 μM Rgg3_St_ and 360 μM SHP3 for 10 min at 25 °C.

EM grids were prepared using a Vitrobot Mark IV autoplunger (FEI), with the environmental chamber at 22 °C and 100% relative humidity. More specifically, 2.5 μL Rgg3_St_-SHP3 was applied to glow-discharged UltraAuFoil (1.2/1.3) 300-mesh grids (Quantifoil), blotted with filter paper for 3-3.5 s, and flash-frozen by plunging in liquid ethane cooled with liquid nitrogen. Grids were then stored in liquid nitrogen.

Cryo-EM data were collected at the Rutgers New Jersey CryoEM/CryoET Core Facility using a 200 kV Talos Arctica (FEI/ThermoFisher) electron microscope equipped with a K2 Summit direct electron detector (Gatan) and a GIF Quantum energy filter (Gatan) with slit width of 20 eV. Data were collected automatically in counting mode using EPU (FEI/ThermoFisher), a nominal magnification of 130,000X, a calibrated pixel size of 1.038 Å per pixel, and a dose rate of 6.347 electrons/pixel/s. Movies were recorded at 200 ms/frame for 8 s (40 frames total), resulting in a total radiation dose of 47.13 electrons/Å^2^. Defocus range was 0.5 to 3.5 μm. A total of 1,464 micrographs were recorded from two grids over two days. Micrographs were gain-normalized and defect corrected.

The processing workflow for the Rgg3_St_-SHP3 dataset is presented is Fig. S1B, with the processing completed entirely within cryoSPARC.v2 (2). Briefly, 1464 micrographs were subjected to patch motion correction and patch CTF estimation. After curation of exposures for estimated resolution, motion distance, motion curvature and ice thickness, 815 micrographs were selected for further processing. Blob picker was used to create templates directly from the data, which were then used with template picker to select 389k particles. These particles were extracted using a 192-pixel box binned to 96 pixels (2.076 Å/pix). The dataset was subjected to multi-class ab initio reconstruction and further 2D classification to yield a 263k particle stack which were re-extracted using an unbinned 256-pixel box. The resulting particles were subjected to two rounds of multi-class ab initio reconstruction with 4 classes, which were subsequently used as starting volumes for heterogeneous refinement. In both cases, the two higher resolution classes were selected for the next round. Subsequently, non-uniform refinement (3) was used, leading to a reconstruction at 3.37 Å from ∼87k particles. Further cleaning of the dataset and a second run of cleanup from the initial stack (after subtracting the first clean particle stack) gave a stack of 129k particles with a 3.25 Å reconstruction. The particles were subjected to local motion correction with minimal effects. After verification that the resulting reconstruction volume is C2 symmetric, C2 symmetry was imposed and used in multi-class ab initio reconstructions to further clean the particle stack reducing noise in the reconstruction. The final stack consisted of ∼44k particles from 428 micrographs, resulted in a 3.04 Å reconstruction after non-uniform refinement. Finally, the particle stack was subjected to global and local CTF refinement and symmetry expansion under C2 symmetry, leading to an effective particle stack of ∼88k and a 2.95 Å reconstruction after local refinement. The final map was sharpened using phenix.auto_sharpen (4).

The initial atomic model for Rgg3_St_-SHP3 was built by first manually aligning and then rigid model refining separately the two Rgg3_St_ protomers from the X-ray crystal structure of Rgg3_St_ alone into the Rgg3_St_-SHP3 cryo-EM map using Coot (5). Subsequent refinement and building of the model was performed using phenix.real_space_refine (6-8) and Coot, respectively. Real-space refinement included global minimization and atomic-displacement parameter (ADP or B-factors) refinement. Restraints included standard geometric, secondary structure, non-crystallographic symmetry (NCS), rotamer, and Ramachandran. Comprehensive model validation was carried out using phenix.validation_cryoem (Table S1). In addition, EMRinger was utilized for model-map validation, yielding an excellent score (Table S1)(9). The 3DFSC server was also used to generate a directional FSC estimate and sphericity measure (10). Molecular graphics for the cryo-EM map were generated using UCSF Chimera and ChimeraX (11, 12).

For the Euler angle distribution plot, the final particles.cs file into a .star file using the UCSF pyem csparc2star.py program and then converting the .star file into a .bild file using the star2bild.py program (Asarnow, D., Palovcak, E., Cheng, Y. UCSF pyem v0.5. Zenodo https://doi.org/10.5281/zenodo.3576630 (2019)). UCSF Chimera was used for visualization of the Euler angle distribution (11).

### Crystallization and Diffraction Data Collection

Crystals of Rgg2_Sd_ in complex with its Rgg box (annealed oligos pSHP2DNA_top and pSHP2DNA_bot) were obtained via the vapor diffusion method by first incubating 86 μM Rgg2_Sd_ with 128 μM Rgg box at room temperature for 10 minutes. After incubation, hanging drops were created using 2 μL of the Rgg2_Sd_-Rgg box mixture combined 1:1 with a mother liquor containing 100 mM trilithium citrate and 24% PEG 3,350 at 20 °C. Prior to X-ray diffraction data collection, crystals were moved to a solution of the mother liquor supplemented with 7% glycerol. X-ray diffraction data for Rgg2_Sd_-Rgg box were collected using single crystals mounted in nylon loops that were then flash-cooled in liquid nitrogen before data collection in a stream of dry N_2_ at 100 K. X-ray diffraction data were collected at the Stanford Synchrotron Radiation Lightsource (SSRL) beamline 14-1 at 1.1808 Å with a MARmosaic 325 CCD detector.

Crystals of Rgg3_St_ in complex with its Rgg box (annealed oligos pSHP3DNA_top and pSHP3DNA_bot) were obtained via the vapor diffusion method by first incubating 133 μM Rgg3_St_ with 133 μM Rgg box and 2.5% dimethyl sulfoxide (DMSO) at room temperature for 10 minutes. After incubation, hanging drops were created using 2 μL of the Rgg3_St_-Rgg box mixture combined 1:1 with mother liquor containing 170 mM ammonium acetate, 85 mM sodium citrate (pH 5.5), 22% PEG 4,000, and 15% glycerol at 20 °C. X-ray diffraction data for Rgg3_St_-Rgg box were collected using single crystals mounted in nylon loops that were then flash-cooled in liquid nitrogen before data collection in a stream of dry N_2_ at 100 K. X-ray diffraction data were collected at SSRL beamline 9-2 at 0.97946 Å with a Dectris Pilatus 6M detector.

Rgg3_St_ crystals were obtained via the vapor diffusion method with 2 uL drops containing 125 μM Rgg3_St_ and 2.5% DMSO combined 1:1 with mother liquor containing 100 mM sodium citrate (pH 5.2), 300 mM Na/K tartrate, and 1.4 M ammonium sulfate. X-ray diffraction data for Rgg3_St_ were collected using single crystals mounted in nylon loops that were then flash-cooled in liquid nitrogen before data collection in a stream of dry N_2_ at 100 K. X-ray diffraction data were collected at SSRL beamline 9-2 at 0.97946 Å with a Dectris Pilatus 6M detector.

X-ray diffraction data for all crystals were processed using HKL3000 (13). Initial crystallographic phases were determined by molecular replacement using Phaser (14) and the previously determined structure of Rgg2_Sd_ (PDB ID 4YV6) (1) as a search model for Rgg2_Sd_-Rgg box and Rgg3_St_-Rgg box. The final models were generated using iterative cycles of model building in Coot (5) and refinement in phenix.refine (6). Initial refinement included simulated annealing as well as rigid body, individual atomic coordinate, and individual B-factor refinement. Later rounds of refinement employed individual atomic coordinate, individual B-factor, and TLS refinement. TLS groups were selected using the TLSMD server (15). During the final rounds of refinement, the stereochemistry and ADP weights were optimized. Insufficient electron density was observed for the following residues in flexible regions of the protein structures, and they were omitted from the model: Rgg2_Sd_-Rgg box residues 1-2 and 277-284 in chain A, and 1-3 and 277-284 in Chain B; Rgg3_St_ residues 1-77 and 101-105; Rgg3_St_-Rgg box residues 1-3 and 284. Ramachandran statistics were calculated in Molprobity (16). Molecular graphics were produced with PyMOL (PyMOL Molecular Graphics System, Version 1.8 Schrödinger, LLC).

### Rgg2_Sd_ and Rgg3_St_ Disulphide Bond Analysis

10 μg of Rgg2_Sd_, Rgg2_Sd_-C45S, Rgg3_St_, and Rgg3_St_-C45S were incubated with SDS loading dye (2% SDS, 10% glycerol, 0.1% bromophenol blue, and 100 mM Tris·HCl [pH 6.8]) with and without 50 mM DTT at room temperature for 10 min. Samples were then heated to 95 °C for 5 min, cooled on ice for 10 min, and then centrifuged at 1,000 × g for 1 min at room temperature. 10 μL of each sample were analyzed using 12% SDS-PAGE at 200 V for 45 min.

### Sequence Alignments

Alignment and similarity calculations were obtained using Clustal Omega and the ESPript 3.0 server using the %Equivalent similarity coloring scheme and a global score of 0.7. Coloring is with respect to identical (red), highly similar (yellow), or below the threshold (white) (17, 18). Aligned sequences include Rgg2 from *S. dysgalactiae* (S. dysg; WP_014612092.1), *S. canis* (WP_003047105.1), *S. agalactiae* (S. agal; AKI95852.1), *S. ictaluri* (S. ictal; WP_008088263.1), and *S. pyogenes* (S. pyog; WP_002990747.1), and Rgg3 from *S. pyogenes* (S. pyog; WP_012560528.1), *S. pneumoniae* (S. pneumo; WP_016397759.1), *S. mitis* (WP_042900990.1), *S. porcinus* (S. porc; WP_143920769.1), and *S. thermophilus* (S. thermo; WP_011225998.1).

### Rgg3_St_ Luciferase Reporter Transcriptional Bioassays

These assays were conducted as previously described (1). Briefly, overnight cultures of GAS were diluted 1:100 into chemically defined medium containing 100 µg/mL spectinomycin. Cultures were incubated at 37° C until reaching an OD_600_ of 0.1, at which time 100 µL of culture was transferred to a 96-well clear-bottomed plate containing indicated concentrations of synthetic SHP3-C8 peptide (DIIIIVGG). A 1% decyl aldehyde solution, the substrate for the luciferase enzyme, suspended in mineral oil, was placed between wells of the microtiter plate. Luminescence and OD_600_ readings were recorded every 5 minutes using a BioTek Synergy2 plate reader. Relative light units (RLUs) were calculated by dividing the luminescence values (counts per second) by the corresponding OD_600_ values. Strain KMT101 served as the host GAS strain for *rgg* orthologs and mutants expressed from plasmid pLZ12-spec. KMT101 was engineered to delete *rgg2* in a strain already containing a deletion of *rgg3* and missense mutations of start codons for both *shp2* and *shp3* (BNL193 *rgg3::cat shp2*_*GGG*_ *shp3*_*GGG*_ (19)).

### Electrophoretic Mobility Shift Assays

The EMSA DNA probe was generated *in vitro* using 5′-FAM-tagged oligonucleotide pair BL63/BL64 (19) in 50 mM Tris and 100 mM NaCl, pH 7.5. DNA probes were annealed in a thermocycler by cooling from 95 °C to 15 °C over 160 minutes and treating with exonuclease I (New England Biolabs) to remove single-stranded DNA. For DNA binding experiments, recombinant purified protein was incubated with 10 nM of the prepared fluorescent probe in 20 mM HEPES buffer pH 7.9, 20 mM KCl, 5 mM MgCl_2_, 0.2 mM EDTA, 0.5 mM CaCl_2_, 10 mg/mL BSA, 12% v/v glycerol, and 0.5 mM DTT for 30 minutes at room temperature. sSHP3-C8 was added to a final volume of 2 µM in indicated reactions; an equal volume of DMSO was added to reactions omitting sSHP3-C8. Samples were loaded onto native polyacrylamide gels buffered with 20 mM potassium phosphate, pH 7.5. Gels were run at 4 °C for 40 minutes at 100 V and fluorescently imaged using a Typhoon phosphorimager (GE Life Sciences).

---

**Fig. S1.**
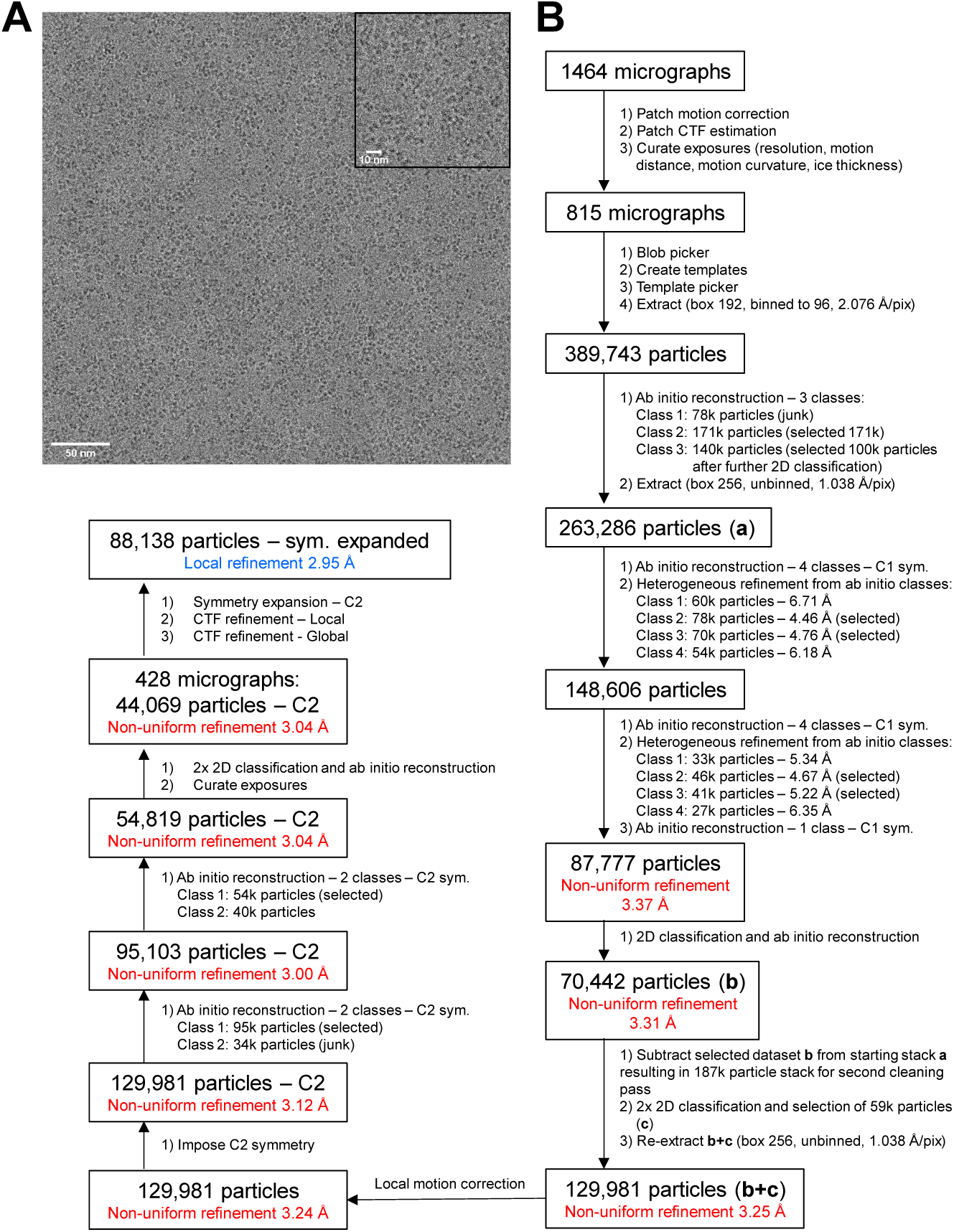
Rgg3_St_-SHP3 cryo-EM structure determination. (A) Representative electron micrograph of Rgg3_St_-SHP3. The scale bar was added with the Fiji implementation of ImageJ (∼4x binned, 0.4154 nm per pixel) (20). The inset shows a ∼2x magnified view of particles from the micrograph. (B) Data processing workflow used to enable high-resolution reconstruction of the Rgg3_St_-SHP3 complex. The workflow is further described in the relevant Supplementary Materials and Methods section.

**Fig. S2.**
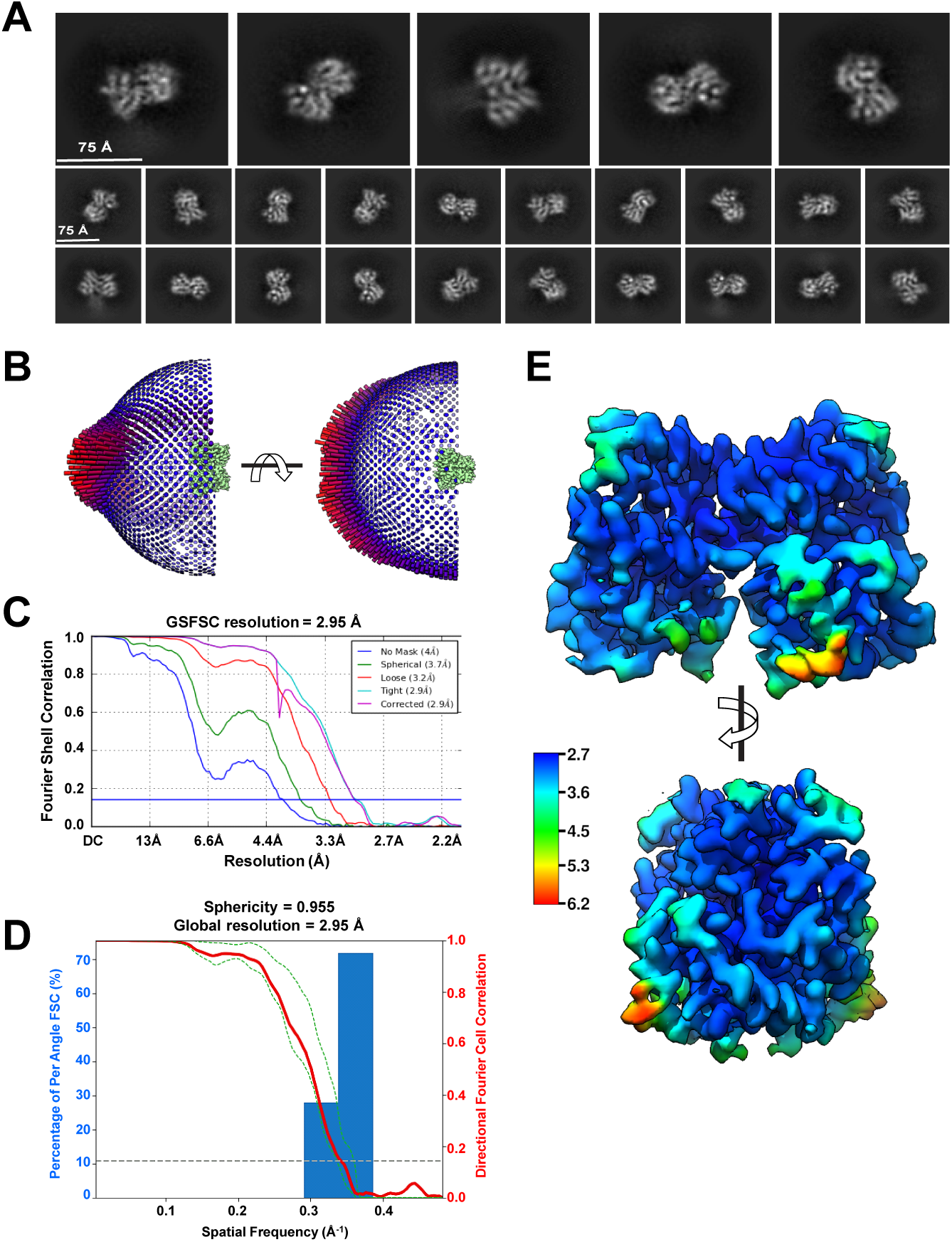
Rgg3_St_-SHP3 cryo-EM data quality and validation. (A) Two-dimensional class averages after reference-free two-dimensional classification of the final particle stack in cryoSPARC.v2 (re-extracted in a 144-pixel box, ∼150 Å). Five majority classes are shown on the top row, with the remainder classes presented below in smaller tiles. (B) Euler angle distribution plot of the final Rgg3_St_-SHP3 particle stack in two directions. See Supplementary Materials and Methods for a description of how this plot was created (C) Fourier shell correlation (FSC) curve for Rgg3_St_-SHP3 generated after the final local refinement in cryoSPARC.v2. (D) Directional FSC curve generated using the 3DFSC server (10). Embedded histograms of directional resolution are shown in blue. Global FSC is shown in red, and ±1 S.D. from the mean of directional FSC is shown as green dashed lines. (E) Local resolution of Rgg3_St_-SHP3 mapped to the final unsharpened reconstruction in two orthogonal views. Local resolution was estimated using an implementation of the blocres program (21) in cryoSPARC.v2 (2). The coloring scale shows a range of resolution from deep blue (2.7 Å) to red (6.2 Å).

**Fig. S3.**
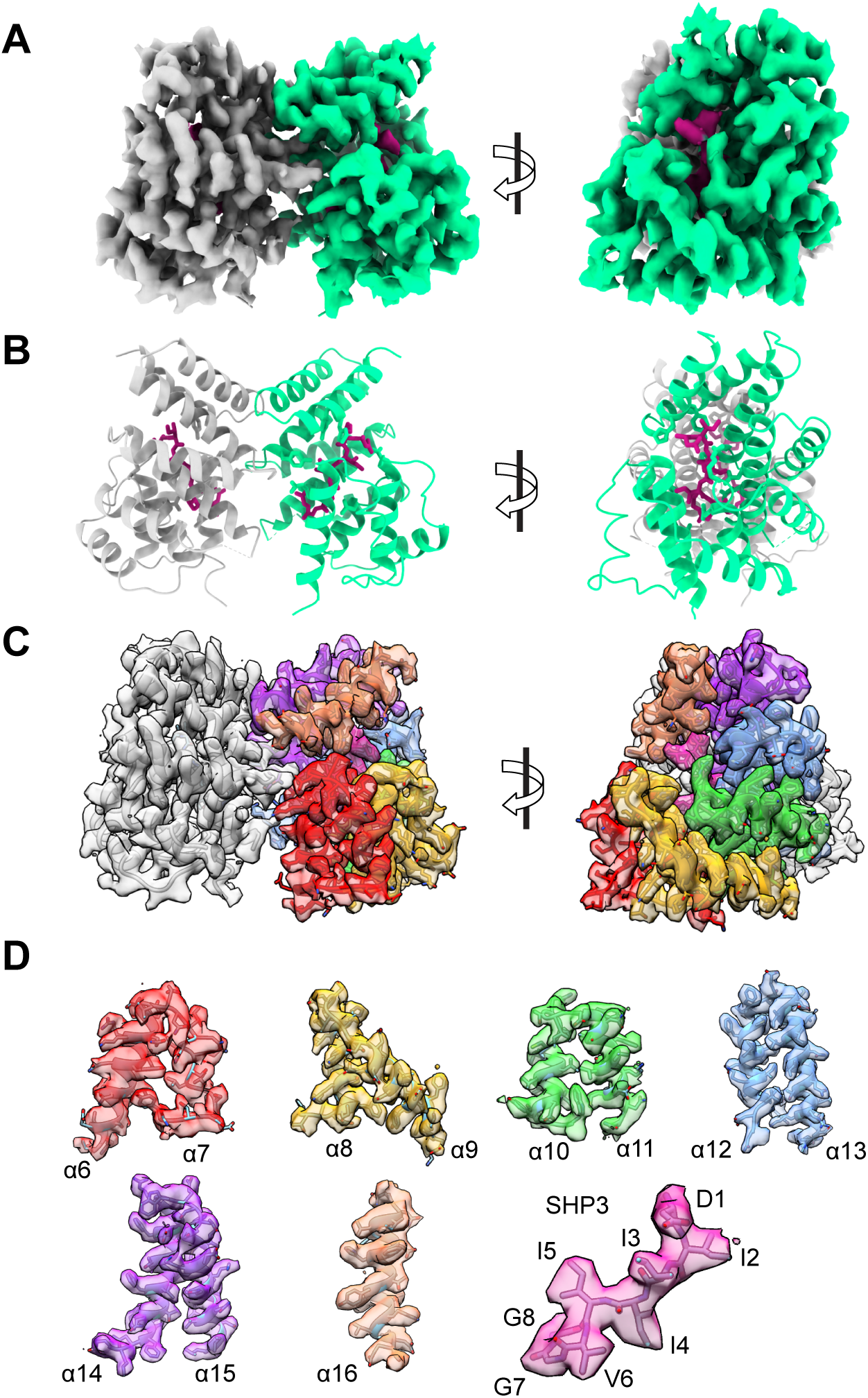
Rgg3_St_-SHP3 cryo-EM map and model. (A) Cryo-EM map of the Rgg3_St_-SHP3 complex at 2.95 Å nominal resolution, contoured at 7.5 σ. Subunit A is shown in green color. The SHP3 peptide is shown in magenta. Two orthogonal views are presented. This representation was generated with Chimera X (12). (B) Ribbon representation of the model corresponding to (A) with a matching color scheme. (C) Superposition of the cryo-EM map and corresponding model of the Rgg3St-SHP3 complex. The map is contoured at 6.5 σ and it is colored according to the color scheme shown in Fig. 1A,B for the repeat domain helices. This representation was generated with Chimera (11). (D) Extracted densities from the cryo-EM to showcase the quality of the map, with corresponding portion of the model superimposed. The map was contoured at 6.5 σ before extracting the partial densities. The density corresponding to the SHP3 peptide from subunit A is shown with magenta coloring and the SHP3 peptide model superimposed. The individual amino acids (DIIIIVGG) are marked in sequence.

**Fig. S4.**
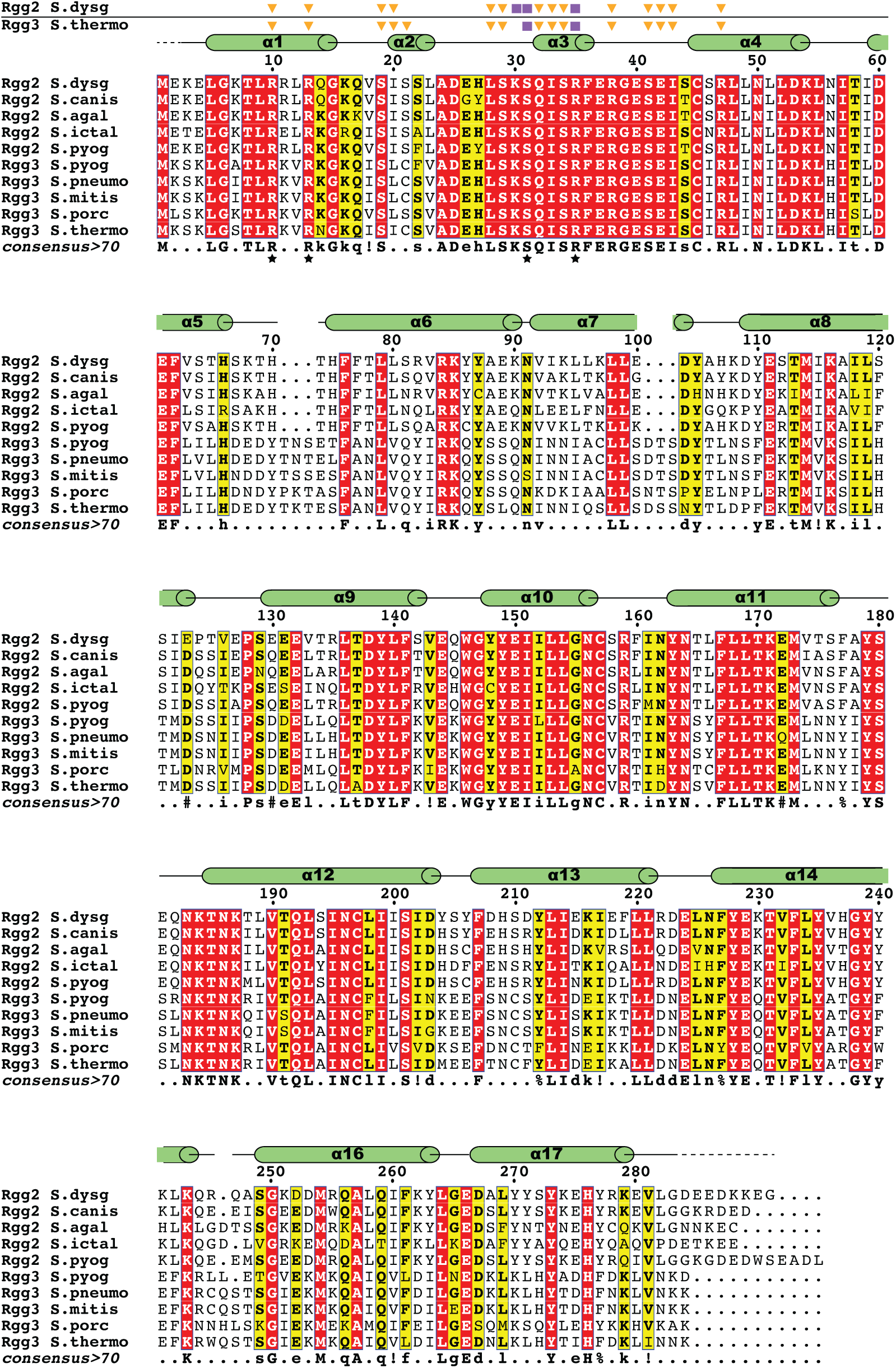
Sequence alignment of representative Rgg2 and Rgg3 proteins. Amino acid numbering and secondary structure elements i.e., α-helices (green cylinders) and loops (black lines), correspond to Rgg3_St_. Rgg2_Sd_ and Rgg3_St_ residues that interact with the Rgg-box phosphate backbone (orange triangles) or nucleotide base (purple squares) are indicated. Residues mutated to alanine and tested in EMSAs are indicated by a black star beneath the consensus sequence and colored as described in Methods.

**Fig. S5.**
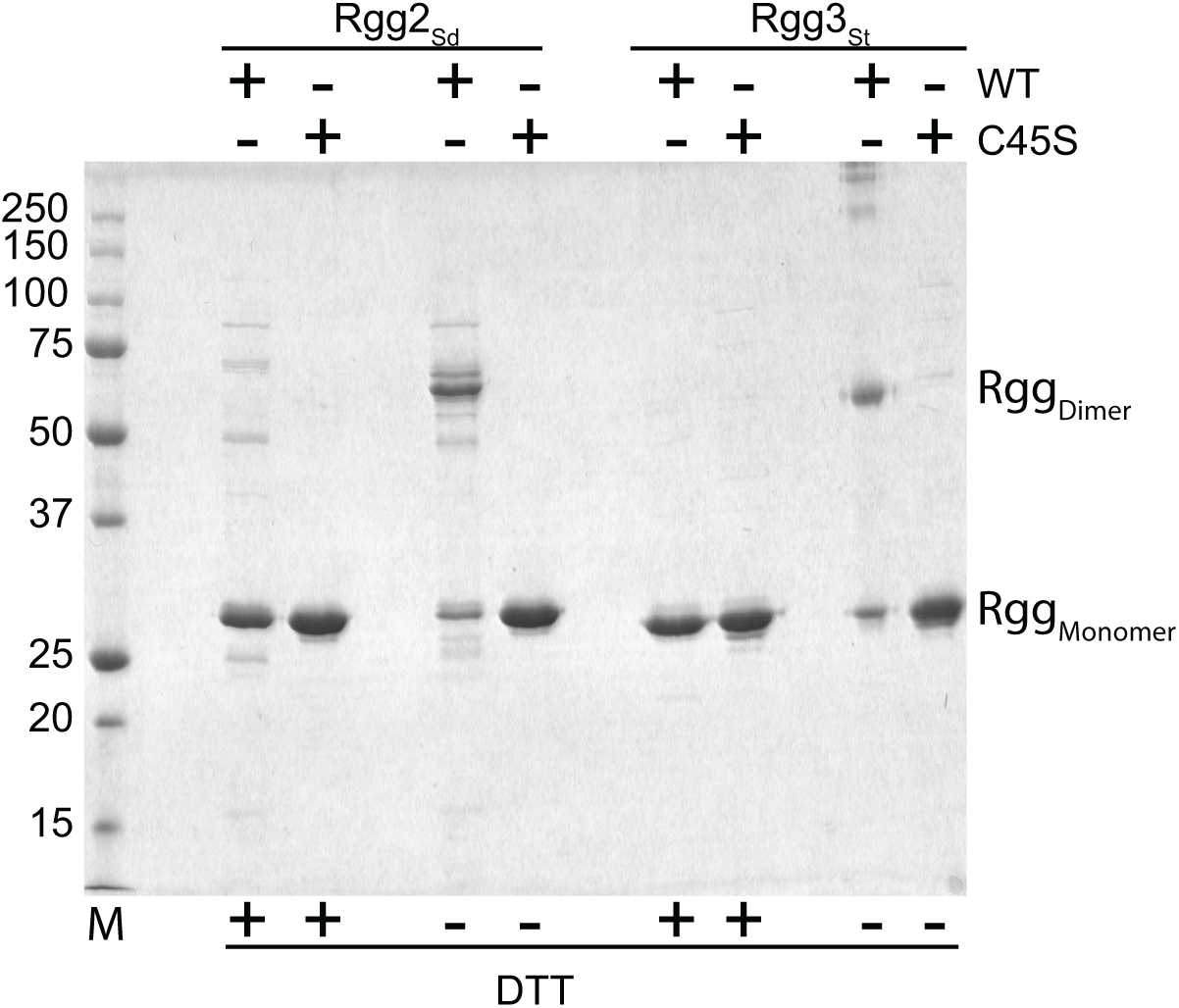
*In vitro* formation of the Rgg3_St_ intermolecular disulphide bond. Samples of Rgg3_St_ were boiled in the presence or absence of DTT before analysis by SDS/PAGE as described in Methods. M, molecular weight standards.

**Fig. S6.**
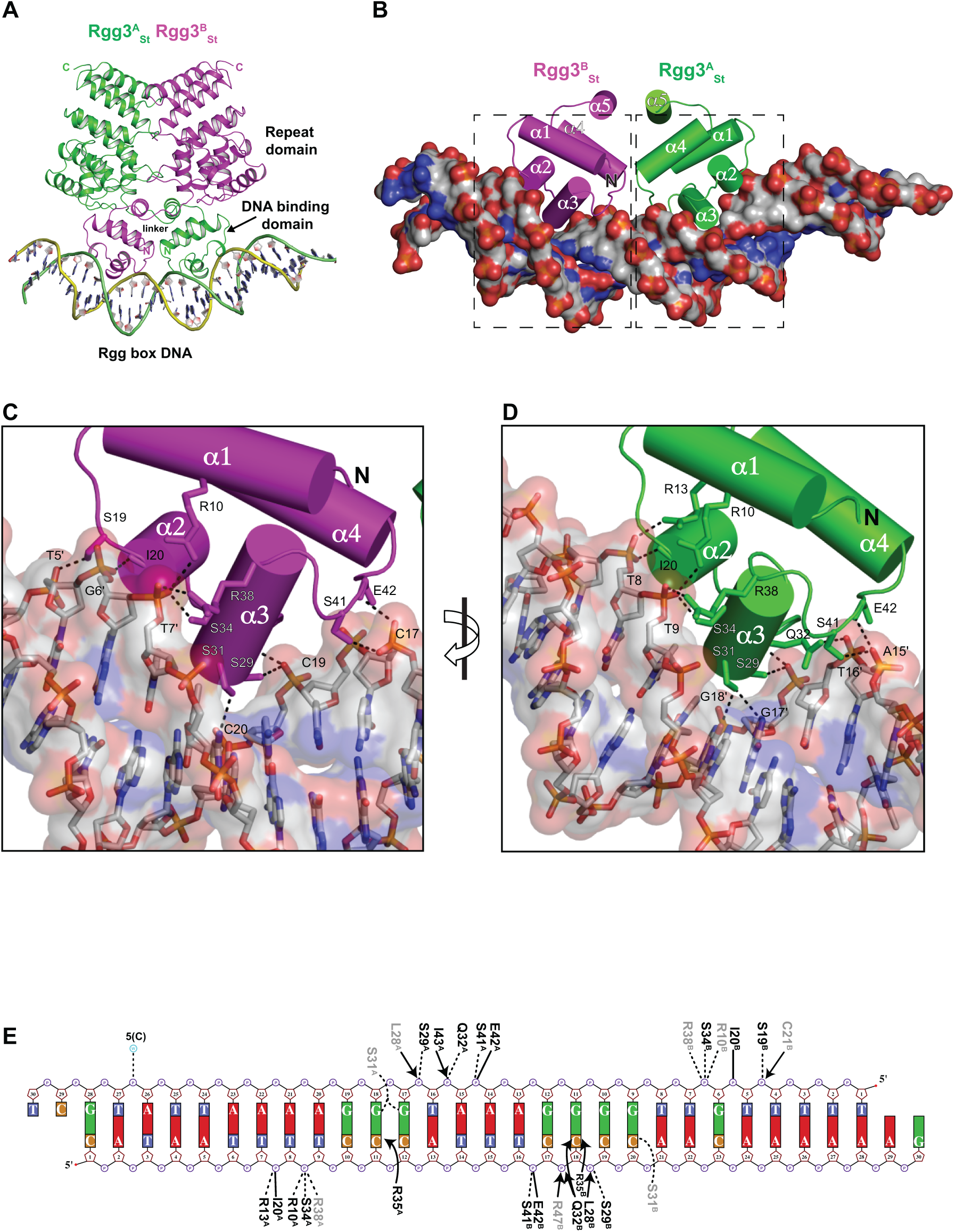
Crystal structure of Rgg3_St_ bound to Rgg3-box DNA. (A) Rgg2_Sd_ dimer bound to Rgg2-box DNA. (B) Isolated view of the Rgg3_St_ DNA binding domains (cylindrical helices) in complex with Rgg-box DNA (surface rendering colored by element). (C and D) Expanded view of the areas enclosed by the dashed black rectangles on the left or right in panel B, respectively. The structures were rotated slightly to provide a clear view of the Rgg4_St_-Rgg box interactions. Rgg-box is depicted as a semi-transparent surface and sticks. Rgg3_St_ side chains or main chains that interact with Rgg-box are labeled and shown as sticks. Hydrogen bonds are depicted as dashed lines. (E) Rgg3_St_-Rgg box interaction schematic. Solid and dashed lines indicate H-bond interactions between Rgg box and the Rgg3_St_ main chain and side chains, respectively. Arrows represent intermolecular hydrophobic interactions (<3.35 Å). Sidechains with ambiguous electron density are shaded grey. DNA phosphate is depicted as circles, ribose sugars as pentagons, and nucleotide bases as rectangles. Portions of the schematic were generated using NUCPLOT (22).

**Fig. S7.**
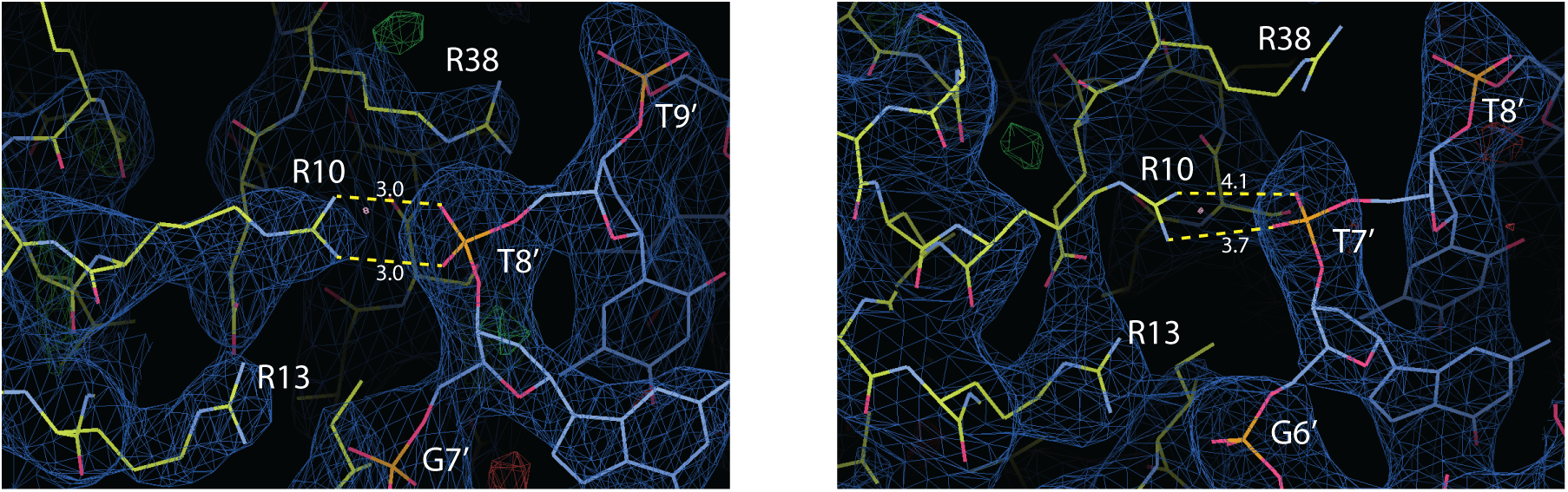
Representative Rgg2_Sd_-Rgg box and Rgg3_St_-Rgg box electron density. Electron density around R10, R13, and R38 in Rgg2^B^_Sd_-Rgg box (left) and Rgg3^B^_St_-Rgg box (right). Protein residues are depicted as yellow sticks, DNA nucleotides as blue sticks. The blue mesh represents the 2F_o_-F_c_ map scaled to 1.0 *σ*. The green and red mesh represent the positive and negative contour F_o_-F_c_ map scaled to 3.0 *σ*, respectively. Dashed lines indicate distances measured in Å. Image generated in Coot (5).

**Fig. S8.**
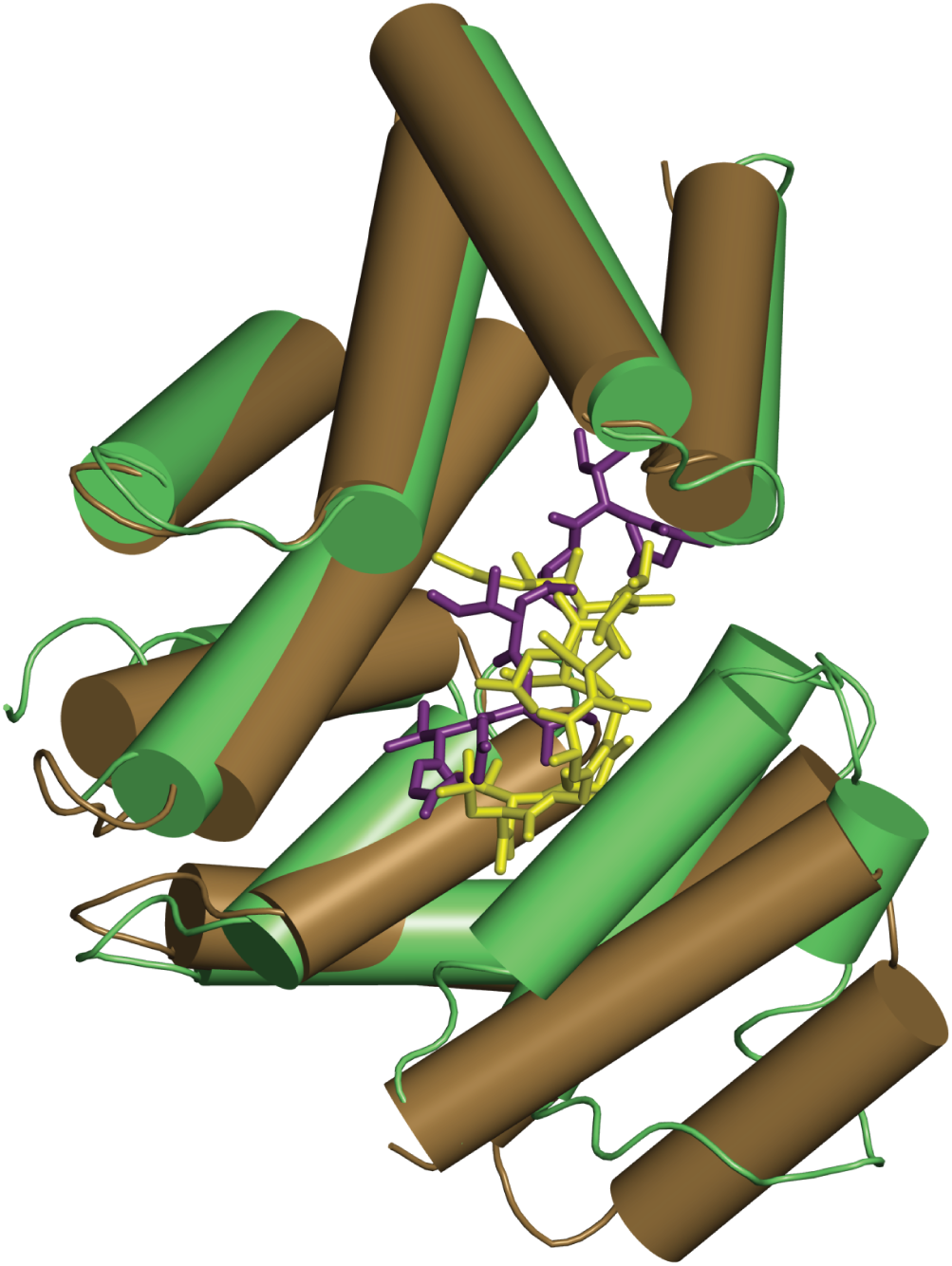
SHP3 and cyclosporin A bind to overlapping sites on their target Rgg proteins. Structural alignment of Rgg3^A^_St_ (green cylinders) bound to SHP3 (yellow sticks), and Rgg2^A^_Sd_ (brown cylinders) bound to CsA^E^ (yellow sticks)(PDB 4YV9. For clarity, the Rgg2^A^_Sd_ DNA binding domain is not shown, and only one monomer of each Rgg2/3 dimer is shown.

**Table S1.**
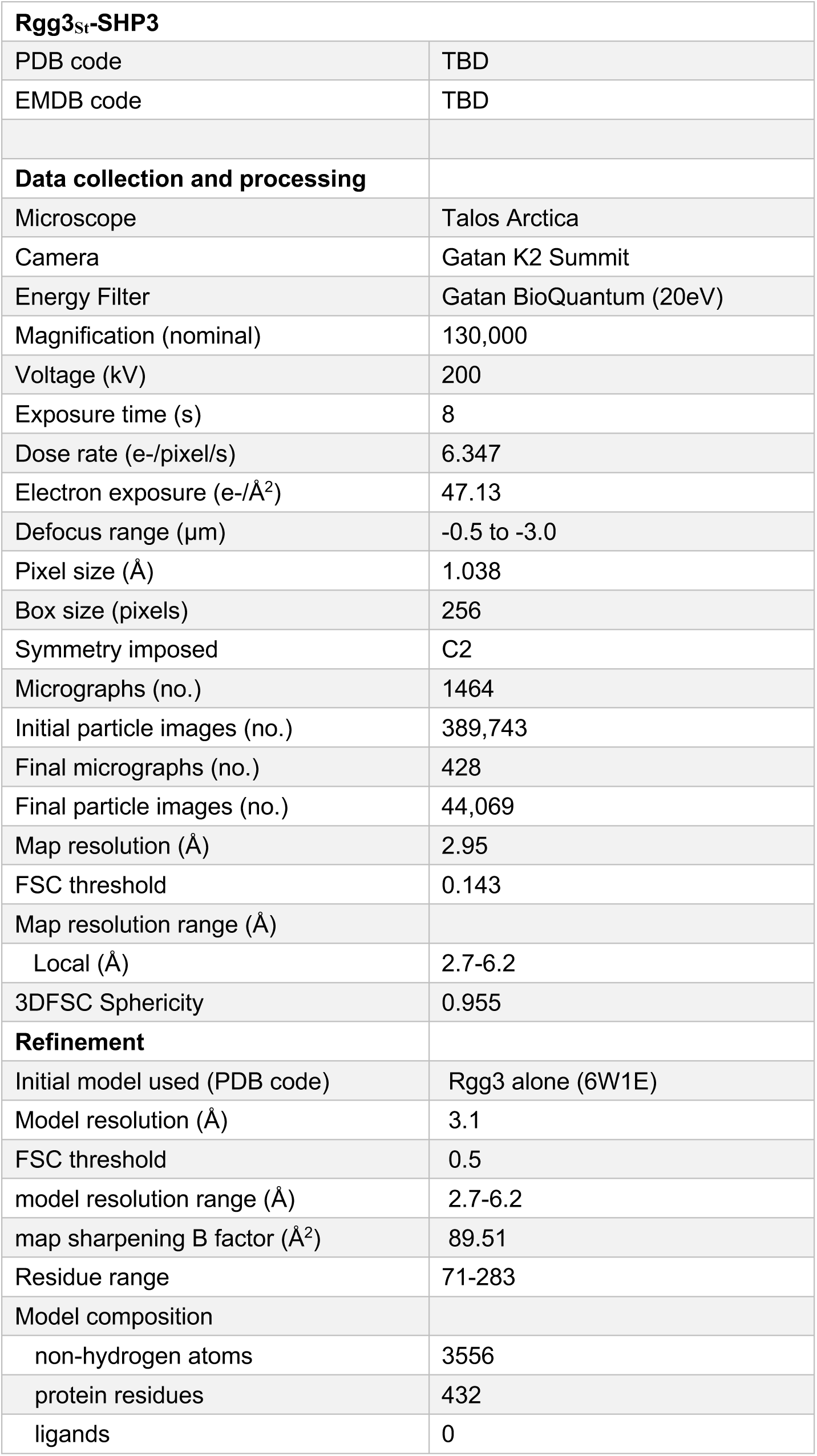
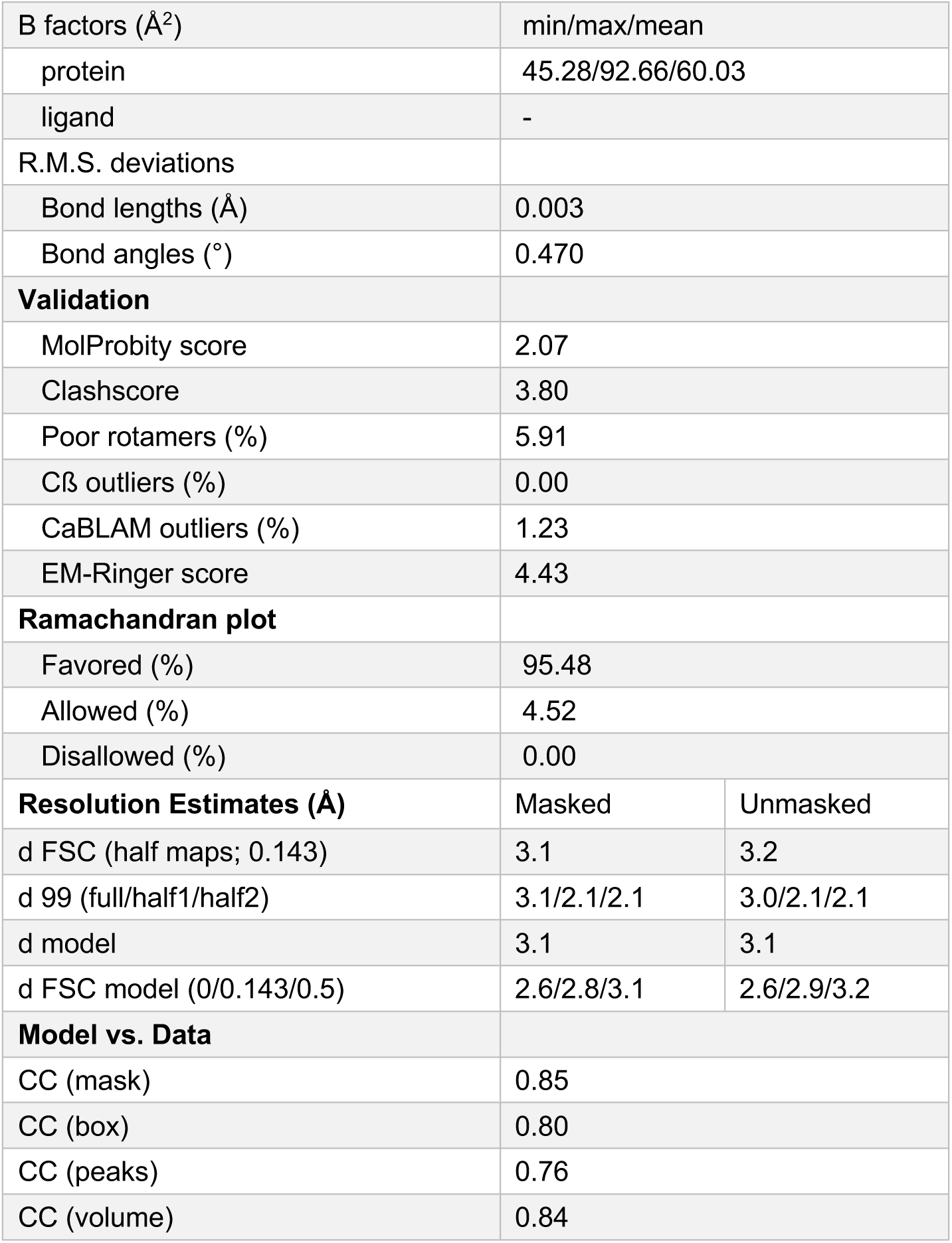
Cryo-EM data collection, refinement, and validation statistics for Rgg3_St_-SHP3 complex structure.

| **Rgg3<sub>St</sub>-SHP3** |  |
| --- | --- |
| PDB code | TBD |
| EMDB code | TBD |
| **Data collection and processing** |  |
| Microscope | Talos Arctica |
| Camera | Gatan K2 Summit |
| Energy Filter | Gatan BioQuantum (20eV) |
| Magnification (nominal) | 130,000 |
| Voltage (kV) | 200 |
| Exposure time (s) | 8 |
| Dose rate (e-/pixel/s) | 6.347 |
| Electron exposure (e-/Å<sup>2</sup>) | 47.13 |
| Defocus range (μm) | -0.5 to -3.0 |
| Pixel size (Å) | 1.038 |
| Box size (pixels) | 256 |
| Symmetry imposed | C2 |
| Micrographs (no.) | 1464 |
| Initial particle images (no.) | 389,743 |
| Final micrographs (no.) | 428 |
| Final particle images (no.) | 44,069 |
| Map resolution (Å) | 2.95 |
| FSC threshold | 0.143 |
| Map resolution range (Å) |  |
| Local (Å) | 2.7-6.2 |
| 3DFSC Sphericity | 0.955 |
| **Refinement** |  |
| Initial model used (PDB code) | Rgg3 alone (6W1E) |
| Model resolution (Å) | 3.1 |
| FSC threshold | 0.5 |
| model resolution range (Å) | 2.7-6.2 |
| map sharpening B factor (Å<sup>2</sup>) | 89.51 |
| Residue range | 71-283 |
| Model composition |  |
| non-hydrogen atoms | 3556 |
| protein residues | 432 |
| ligands | 0 |

|  |  |  |
| --- | --- | --- |
| B factors (Å<sup>2</sup>) | min/max/mean |  |
| protein | 45.28/92.66/60.03 |  |
| ligand | - |  |
| R.M.S. deviations |  |  |
| Bond lengths (Å) | 0.003 |  |
| Bond angles (°) | 0.470 |  |
| **Validation** |  |  |
| MolProbity score | 2.07 |  |
| Clashscore | 3.80 |  |
| Poor rotamers (%) | 5.91 |  |
| Cβ outliers (%) | 0.00 |  |
| CaBLAM outliers (%) | 1.23 |  |
| EM-Ringer score | 4.43 |  |
| **Ramachandran plot** |  |  |
| Favored (%) | 95.48 |  |
| Allowed (%) | 4.52 |  |
| Disallowed (%) | 0.00 |  |
| **Resolution Estimates (Å)** | Masked | Unmasked |
| d FSC (half maps; 0.143) | 3.1 | 3.2 |
| d 99 (full/half1/half2) | 3.1/2.1/2.1 | 3.0/2.1/2.1 |
| d model | 3.1 | 3.1 |
| d FSC model (0/0.143/0.5) | 2.6/2.8/3.1 | 2.6/2.9/3.2 |
| **Model vs. Data** |  |  |
| CC (mask) | 0.85 |  |
| CC (box) | 0.80 |  |
| CC (peaks) | 0.76 |  |
| CC (volume) | 0.84 |  |

**Table S2.** Statistics for X-ray crystallographic data collection and refinement.

| **PDB ID** | Rgg2<sub>sd</sub>-Rgg box<br>6W1A | Rgg3<sub>st</sub><br>6W1E | Rgg3<sub>st</sub>-Rgg box<br>6W1F |
| --- | --- | --- | --- |
| **Data collection** |  |  |  |
| Space group | P6<sub>5</sub> | P3<sub>2</sub>21 | P6<sub>5</sub> |
| Cell dimensions<br>a, b, c (Å) | 110.27, 110.27,<br>174.00 | 110.36, 110.36,<br>54.93 | 90.79, 90.79, 245.10 |
| $\alpha, \beta, \gamma$ (°) | 90.0, 90.0, 120.0 | 90.0, 90.0, 120.0 | 90.0, 90.0, 120.0 |
| Resolution (Å) | 50.00-2.80 (2.90-2.80) | 50.00-2.20 (2.26-2.20) | 50.00-3.20 (3.29-3.20) |
| Wavelength (Å) | 1.18076 | 0.97946 | 0.97946 |
| Completeness (%) | 99.9 (99.7) | 97.7 (86.7) | 99.5 (100.0) |
| R<sub>sym</sub> (%) | 10.2 (151.8) | 8.1 (47.3) | 5.4 (66.4) |
| CC<sup>1/2</sup> | 0.999 (0.648) | 0.998 (0.939) | 0.986 (0.896) |
| Average I / σI | 31.1 (1.8) | 42.6 (2.3) | 35.4 (2.5) |
| Redundancy | 8.3 | 9.4 | 10.3 |
| Total reflections | 245,818 | 181,920 | 193,899 |
| Unique reflections | 29,533 | 19,380 | 18,729 |
| **Refinement** |  |  |  |
| R<sub>work</sub> / R<sub>free</sub> (%) | 23.35 (35.17) / 26.43<br>(34.82) | 26.51 (34.41) / 30.86<br>(37.74) | 21.68 (29.27) / 26.62<br>(40.68) |
| **Number of atoms** |  |  |  |
| All atoms | 5,865 | 1,712 | 5,842 |
| Protein | 4,588 | 1,678 | 4,606 |
| DNA | 1,230 | 0 | 1,230 |
| Ligands | 28 | 10 | 0 |
| Water | 19 | 24 | 6 |
| **Average B-factor**<br>**(Å<sup>2</sup>)** |  |  |  |
| All atoms | 99.03 | 78.66 | 151.78 |
| Protein | 99.91 | 78.47 | 130.49 |
| DNA | 125.61 | n.d | 231.74 |
| Ligands | 119.31 | 131.94 | n.d. |
| Water | 68.65 | 69.38 | 110.45 |
| **R.m.s. deviations** |  |  |  |
| Bond lengths (Å) | 0.004 | 0.002 | 0.004 |
| Bond angles (°) | 0.811 | 0.518 | 0.779 |
| **Ramachandran**<br>**statistics** |  |  |  |
| Favored (%) | 95.40 | 95.96 | 93.34 |
| Allowed (%) | 4.42 | 3.54 | 6.12 |
| Outliers (%) | 0.18 | 0.50 | 0.54 |

Data collection and refinement statistics. $R_{\text{sym}} = \sum_h \sum_i |I_i(h) - \langle I(h) \rangle| / \sum_h \sum_i I_i(h)$ , where $I_i(h)$ is the $i^{\text{th}}$ measurement of $h$ and $\langle I(h) \rangle$ is the mean of all measurements of $I(h)$ for reflection $h$ . $R_{\text{work}} = \sum ||F_o| - |F_c|| / \sum |F_o|$ , calculated with a working set of reflections. $R_{\text{free}}$ is $R_{\text{work}}$ calculated with only the test set of reflections. Data for the highest resolution shell are given in parentheses. The structures were determined using single crystals.

**Table S3.**
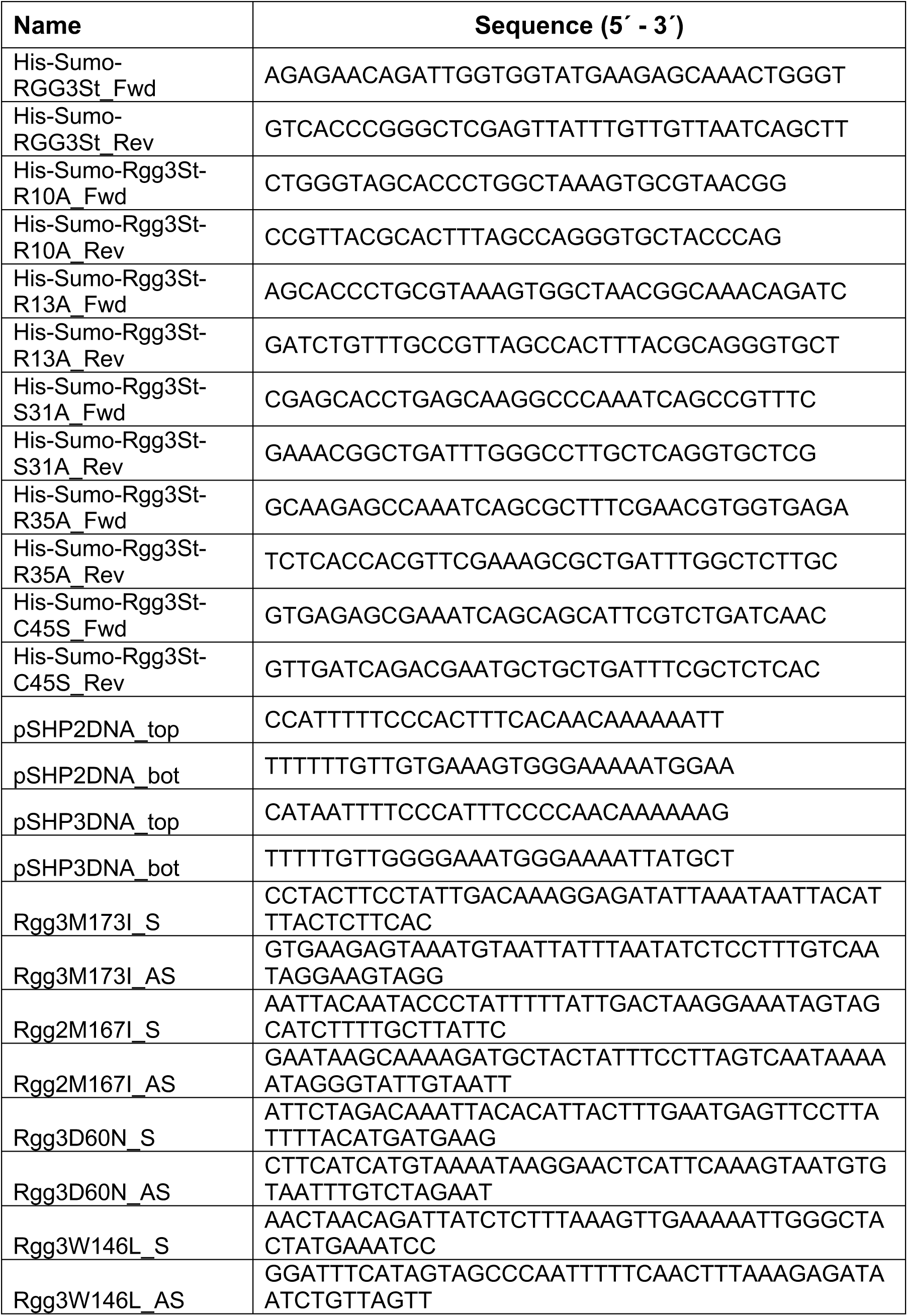
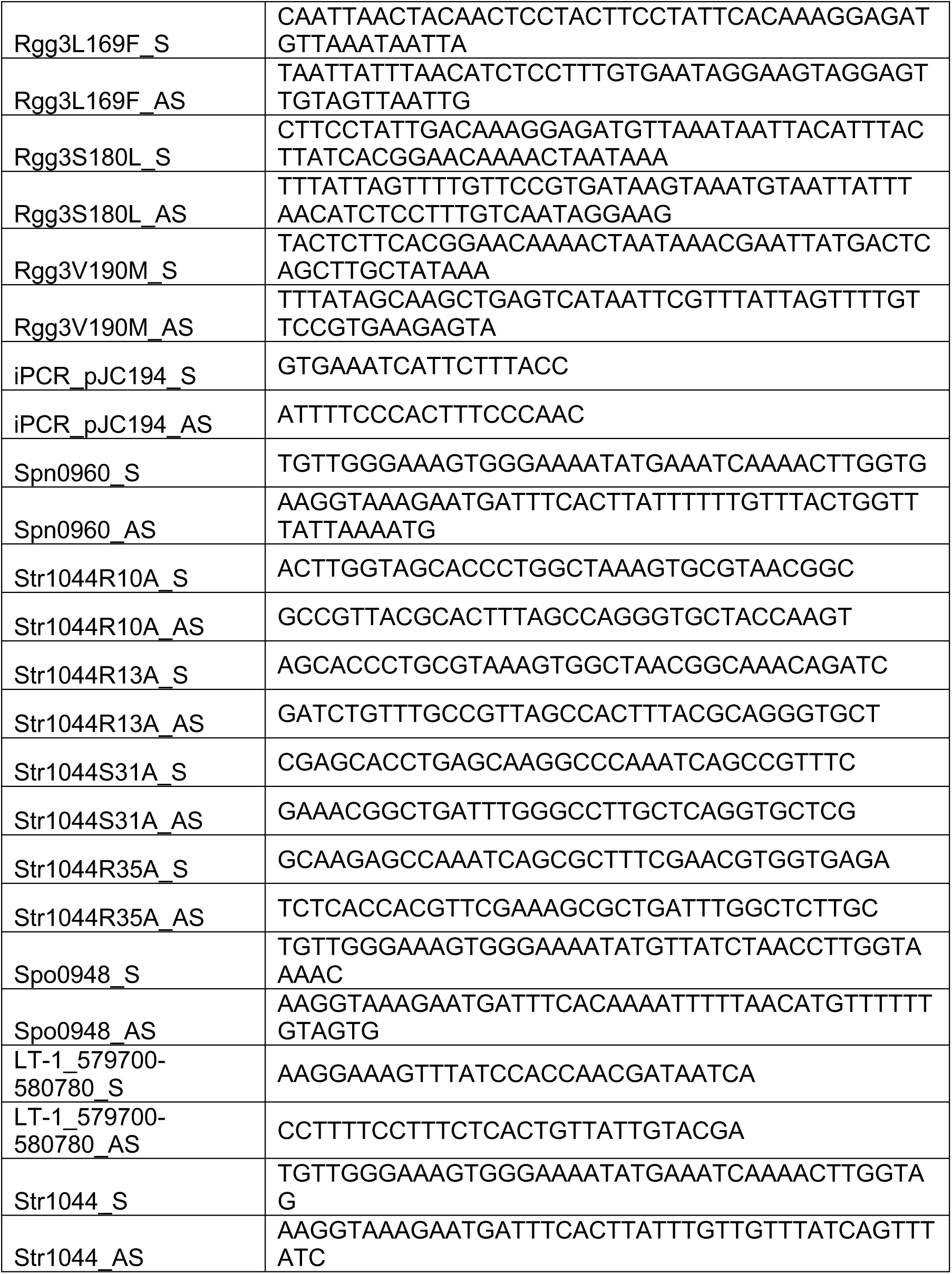
Primers used in this study.

